# Strong experimental support for the hologenome hypothesis revealed from *Drosophila melanogaster* selection lines

**DOI:** 10.1101/2021.09.09.459587

**Authors:** Torsten Nygaard Kristensen, Anna A. Schönherz, Palle Duun Rohde, Jesper Givskov Sørensen, Volker Loeschcke

**Author notes:** Corresponding author: Anna A. Schönherz, Blichers Allé 20, building D23, Tjele, DK 8830, Denmark, e mail. Shared first authorship (Torsten Nygaard Kristensen, Anna A. Schönherz).

## Abstract

Recently it has been proposed, that the holobiont, i.e., the host and its associated microbiome, constitute a distinct biological entity, on which selection operates. This is a fascinating idea that so far has limited empirical justification. Here *Drosophila melanogaster* lines from a large-scale artificial selection experiment, where we selected for stress resistance traits and for longevity, were used to test the hologenome hypothesis. We raised flies from all selection regimes, including a regime where flies were kept at benign standard laboratory condition (control regime) throughout the duration of the experiment, under common garden conditions and sequenced the microbiome of the flies. We found abundant differences in microbial communities between control and selection regimes, but not between replicate lines within the regimes, and microbial diversity was higher in selected relative to control lines. Several major core *Drosophila* bacterial species were differentially abundant in the different selection regimes despite flies being exposed to similar nutritional and general environmental conditions. Our results support the idea that the host and microbiome genomes have evolved in concert and provide experimental support for the hologenome theory of evolution.

## Introduction

Genetic variation is a prerequisite for evolution to occur and selection on standing genetic variation can lead to marked changes in genotypes and phenotypes; thus, selection is a strong driver for evolutionary changes within species and partly governs the process of speciation. The field of evolutionary biology dating back to Darwin’s and Mendel’s revolutionary discoveries and the modern synthesis are important for almost all biological fields including animal and plant breeding, evolutionary adaptation, conservation genetics and for understanding human disease etiologies [1–3]. New tools in molecular biology allow for assessment of endophenotypes such as the transcriptome, the proteome and the metabolome but also for sequencing not only host organisms but also the myriad of microorganisms that coexist with these hosts [4,5]. The simultaneous study of hosts and their microorganisms is a fast developing and novel research field that opens up for investigating the importance of microbial diversity and how the abundance and distribution of different microbes affect the host’s fitness and whether evolutionary forces shape the host and its microbes uniformly [6].

Numerous studies have documented that the microbiome has strong impact on host fitness components including lifespan, fecundity, immune responses, disease resistance, growth or development, and environmental stress tolerance [7–14]. It has also been revealed, that the host can exert some control of the microbial composition, e.g. through nutrient availability by diet choice or host metabolism [15–17], by immune factors [14], or mechanical control such as gut peristalsis [18]. For example, high abundance of certain species of *Lactobacillus* or *Acetobacter* in nutrient poor environments can critically increase *Drosophila melanogaster* larvae growth so that growth levels reach those normally observed on high-protein diets [19,20], suggesting a rescue effect offered by these microbes. Thus, interactions between the host genotype and the microbiota can have strong consequences for the host [21] but the impact of these interactions on evolutionary changes in natural populations is currently poorly understood [22,23].

The growing realization that interactions between microbes and their hosts shape many aspects of life is fundamentally changing the way we think about many fields of biology including evolutionary genetics, disease etiology, and adaptation to changing environmental conditions [24–27]. The ‘holobiont’ concept describes the idea that eukaryote individuals do not act as autonomous units, but rather as networks consisting of the host and all its associated microorganisms. Their collective genomes, the ‘hologenome’, may form a single unit of selection and it has been proposed that this hologenome can evolve [28,29]. For instance, it has recently been shown that selection for increased cold tolerance in the blue tilapia (*Oreochromis aureus*) led to significant alteration of the microbial composition and modulated the microbes’ response to temperature stress in this tropical fish [30].

Likewise, a study investigating symbiosis between the aphid *Acyrthosiphon pisum* and the bacterium *Buchnera aphidicola*, has revealed how a single nucleotide deletion in the bacterium affects the heat-shock transcriptional promoter for ibpA, encoding a small heat-shock protein and that this mutation governs thermal tolerance of the aphid hosts [31]. As a last example providing support for the holobiont theory we have found that the resident microbes in *D. melanogaster* can respond to thermal acclimation, and subsequently contribute to improve the hosts survival to more extreme thermal conditions [13]. Studies like these suggest that the microbiome and symbiont microevolution have strong importance for host’s ability to cope with variable and periodically stressful environmental conditions and thereby for the evolutionary adaptation to such challenges and they support the idea that host and microbiomes may evolve in concert.

Genomic studies aiming at pinpointing signatures of selection often report little evidence for selection at the genomic or endophenotypic level despite strong directional selection and apparent phenotypic responses to selection [32]. One reason for this is that most variation in quantitative genetic traits is influenced by a very large number of genes each contributing with a small effect on the trait in question [33]. Identifying these genes, transcripts or proteins of small effect is a challenge from a statistical point of view [34]. However, an overlooked reason for apparent lack of strong signatures of selection may be that only part of the hologenome, the host genome, is traditionally being investigated in these studies. Evolution of the myriad of microbial genomes impacted by selection may explain a significant proportion of the response in host phenotypes not captured by analysis of changes in host genomes.

Current knowledge suggests that the microbial composition of the host is important for multiple fitness components. It is also clear that the host exerts some control over the microbiota, that the host and microbiota genomes interact on host fitness, leading to the idea that the combined hologenome may evolve and dictate evolutionary changes [35]. This knowledge provides a background for testing how directional selection for host fitness components has affected the distribution and abundance of associated microbes. This is highly relevant to investigate because we have little knowledge on how artificial selection for specific host traits impacts and may be facilitated by evolution of the microbiome. We propose that interactions between host and microbial genomes govern evolutionary responses. This is investigated using *D. melanogaster* lines from a highly replicated artificial selection experiment. We do this by sequencing the microbiome of populations of *D. melanogaster* selected for different stress resistance traits and longevity and from their corresponding non-selected control populations [36,37]. Lines from each selection regime and lines that had not experienced artificial directional selection, i.e., the unselected control lines, were all reared at similar benign environmental conditions, and they subsequently had their microbiome sequenced. This unique material allows us to test the hypothesis that directional selection for numerous fitness components in *D. melanogaster*, leads to marked and selection-regime-specific changes in microbial composition. We find strong support that the microbiome has evolved alongside the host and that a mutualistic relationship between host and microbiome partly explains the evolved phenotypic characteristics of the host. These results constitute convincing support for the hologenome hypothesis of evolution and suggest that adaptation to stressful environmental conditions critically depends on interactions between host and microbial genomes.

## Material and Methods

### Description of selection regimes and original experimental setup

The flies used in the current study originated from a large environmental-stress and life-history trait selection experiment described previously [36]. In short, initially a mass-bred laboratory population of *D. melanogaster* was established and maintained at 25 °C in 12-hour light-dark cycles, on standard oatmeal-sugar-yeast-agar medium. Six artificial selection regimes were established together with one unselected control (UC) line. For each selection regime (and control) five independent replicated lines were generated. In each selection regime there was selected for one of the following traits: increased desiccation resistance (DS), increased longevity (LS), increased cold-shock resistance (CS), increased heat-shock resistance (HS), increased heat knockdown resistance (KS) and increased starvation resistance (SS). The selection was applied every second generation to allow for recovery and to avoid transgenerational effects [38,39].

The initial phenotypic assessment was performed after 21 generations of selection (45 generations in total), except for LS and SS which were selected for 11 and 26 generations, respectively [36]. Flies from each of the selection regimes (and the unselected controls) have previously been characterized at the genomic level [40], and on a range of different endophenotypic levels [41–44]. The flies used in the current study were collected after 32 generations of selection (i.e., 67 generations of maintenance) for the HS, CS, KS, DS and UC regimes, whereas flies from the SS lines were from generation 81 (i.e., 37 selection events), and flies from the LS regime were from generation 31 (i.e., 14 generations of selection). Phenotypic assessments of selection responses were not performed in the generation where flies were harvested for microbiome analysis and therefore the responsiveness to selection is based on results from the data provided in Bubliy and Loeschcke [36].

### Microbiome community analysis by 16S rRNA quantification

A total of 20 females were sampled from each of the biological replicates of the selection and control lines, snap frozen in liquid nitrogen, and stored at −80 °C for 14 years before performing the microbiome analysis reported here.

DNA extraction and sequencing of 16S rRNA were conducted externally by DNASense ApS (Aalborg, Denmark, https://dnasense.com/). DNA was extracted from pools of 20 female flies using DNeasy Blood and Tissue kit following the manufacturer’s recommendations (Qiagen, Germany). Bacterial 16S V1-V3 rRNA libraries were prepared using a custom protocol based on Caporaso et al. (2012) and amplified using V1-V3 specific primers: [27F] AGAGTTTGATCCTGGCTCAG and [534R] ATTACCGCGGCTGCTGG [45]. The amplicon libraries were purified using a standard protocol for Agencourt Ampure XP Beads (Beckman Coulter, USA) with a bead to sample ratio of 4:5. Purified libraries were pooled in equimolar concentrations and diluted to 6 nM. Samples were paired-end sequenced (2×300 bp) on a MiSeq platform (Illumina, USA) using the MiSeq Reagent kit v3 (Illumina, USA) according to standard guidelines.

### Bioinformatic processing and analysis

Demultiplexed fastq files were processed with QIIME2 v2019.10 [46]. Forward and reverse primers were removed using the cutadapt plugin v2019.10 [47]. To infer amplicon sequence variants (ASVs), trimmed reads were quality filtered, denoised, merged, and PCA chimeras were removed using the DADA2 plugin v2019.10 [48]. For quality filtration, paired-end sequences were truncated at 297 and 261 bp for forward and reverse reads, respectively (50-percentile Phred score quality drop below 30 and 28, respectively). DADA2 default parameter settings were applied otherwise. A total of 304 ASVs were detected. ASVs were taxonomically assigned using a 16S V1-V3 specific Naive Bayes classifier [49]. The classifier was trained on 99% similarity clustered 16S rRNA gene sequences extracted from the SILVA v132 reference database [50], and trimmed to only include the V1-V3 region bound by the 27F/534R primer pair. For phylogenetic inference, ASV sequences were aligned with Mafft v7.310 [51], highly variable positions were masked, an unrooted phylogenetic tree was constructed with FastTree v2.1.10 [52], and the tree was rooted by the midpoint of the longest tip-to-tip distance in QIIME2.

Subsequent microbial data analyses were performed with R v3.6.2 [53]. ASVs unassigned at phylum level and ASVs assigned as chloroplasts, mitochondria, or cyanobacteria were removed from further analyses (303 ASVs remaining). Rarefaction curves were generated with the vegan package v2.5-6 [54] and uneven sampling depth was scaled to minimum read depth by rarefying data to the minimum number of reads across all samples (13.723 sequence reads per sample; 297 ASVs remaining) using the *rarefy_even_depth* function implemented in the phyloseq package v1.30.0 [55]. Overall microbial composition was investigated in rarefied data. Top 10 most abundant bacterial genera across, as well as within, selection regimes were identified and visualized as heatmaps using the ampvis2 package v2.5 [56]. ASV counts were converted to within-sample relative abundances and for each sample, microbial compositions were visualized at phylum and family level using the microbiome package v1.8.0 [57]. For subsequent diversity comparisons, the rarefied count data were pruned based on ASV prevalence, removing ASVs present in less than two samples and with total abundance less than 0.01% across all samples resulting in 141 remaining ASVs (99.4% of rarefied and 48.3% of processed sequence reads). If not stated directly, subsequent analyses were conducted for filtered, rarefied and pruned ASV data. Inter-sample complexity (alpha diversity) was investigated using ASV count data, whereas for intra-sample diversity (beta diversity) ASV count data were converted to relative abundances.

### Inter-sample complexity

Alpha diversity estimates including the observed number of ASVs (richness) and the Shannon’s diversity index (considering richness and evenness of ASV abundances) were computed with phyloseq. Faith’s phylogenetic diversity (PD) index, a measure of diversity which incorporates phylogenetic diversity between ASVs, was estimated with the picante package v1.8.2 [58]. Normality of data was tested using the Shapiro-Wilk test. Overall differences in alpha diversity between selection regimes were tested using a one-way ANOVA followed by Tukey’s multiple comparison of means test for pairwise comparisons of selection lines.

### Intra-sample complexity

Beta diversity of the selection and control lines was investigated by Principal Coordinate Analysis (PCoA) using Bray–Curtis dissimilarity distances. Bray-Curtis dissimilarity distances among samples were estimated using phyloseq. PCoA ordination plots were generated with the ggplot2 package v3.3.1 [59]. To determine differences in microbial diversity between selection regimes a permutational multivariate analysis of variance (PERMANOVA) was computed for Bray-Curtis distances using the *adonis* function implemented in vegan, applying 999 permutations and default settings otherwise. Homogeneity of group dispersions (variance) was verified using the *betadisper* function implemented in vegan. *P*-values ≤ 0.05 were considered statistically significant. In addition, a hierarchical cluster analysis for Bray–Curtis dissimilarity distances was computed using the *hclust* function implemented in the stats package [53] applying default settings and visualized as dendrogram using ggplot2.

### Differential abundance analysis

Differential ASV abundances between selection regimes were determined at genus level using a negative binomial generalized linear model approach implemented in the DESeq2 package v1.2.6. [60]. Briefly, taxonomic assignments were agglomerated to genus level using the *tax_glom* function implemented in phyloseq. ASV counts were normalized applying the variance-stabilizing transformation approach implemented in DESeq2, returning log2 scale transformed data. Size factors for each ASV were estimated applying a median-ratio-method. Dispersions of ASV counts were computed and a negative binomial WaldTest was performed [60,61]. For ASV with zero counts a pseudo-count of one was added [62]. Pairwise comparisons were computed for the following six contrasts: CS vs. UC, DS vs. UC, HS vs. UC, KS vs. UC, LS vs. UC, and SS vs. UC. To correct for multiple hypothesis testing, *P*-values were adjusted using the Benjamini-Hochberg procedure [63]. ASVs were considered differentially abundant for adjusted *P*-values ≤ 0.01 and an FDR cut-off of 5%. Results were visualized using ggplot2.

## Results

Illumina 16S rRNA amplicon sequencing returned 1,538,911 raw sequence reads across all 35 samples, ranging from 21,712 to 70,668 reads per sample (mean 43,969). After pre-processing with Qiime2 988,415 sequencing reads remained, representing 64.27% of the raw sequence reads on average. Pre-processed reads detected per sample ranged from 13,723 to 52,185 (mean 28,240). Model based inference of amplicon sequence variants revealed 304 bacterial amplicon sequence variants (ASVs) across all samples, ranging from 12 to 110 unique ASVs detected per sample (mean 59). A total of 141 bacterial ASVs passed filtration procedures (per-sample range: 9 to 86 ASVs; mean: 48 ASVs).

### Microbial Composition

Detected ASVs were assigned to four phyla: Firmicutes (*N*=228), Actinobacteria (*N*=34), Proteobacteria (*N*=32), and Bacteroidetes (*N*=3). Relative abundances at phylum and family level are summarized in Figure 1.

**Figure 1.**
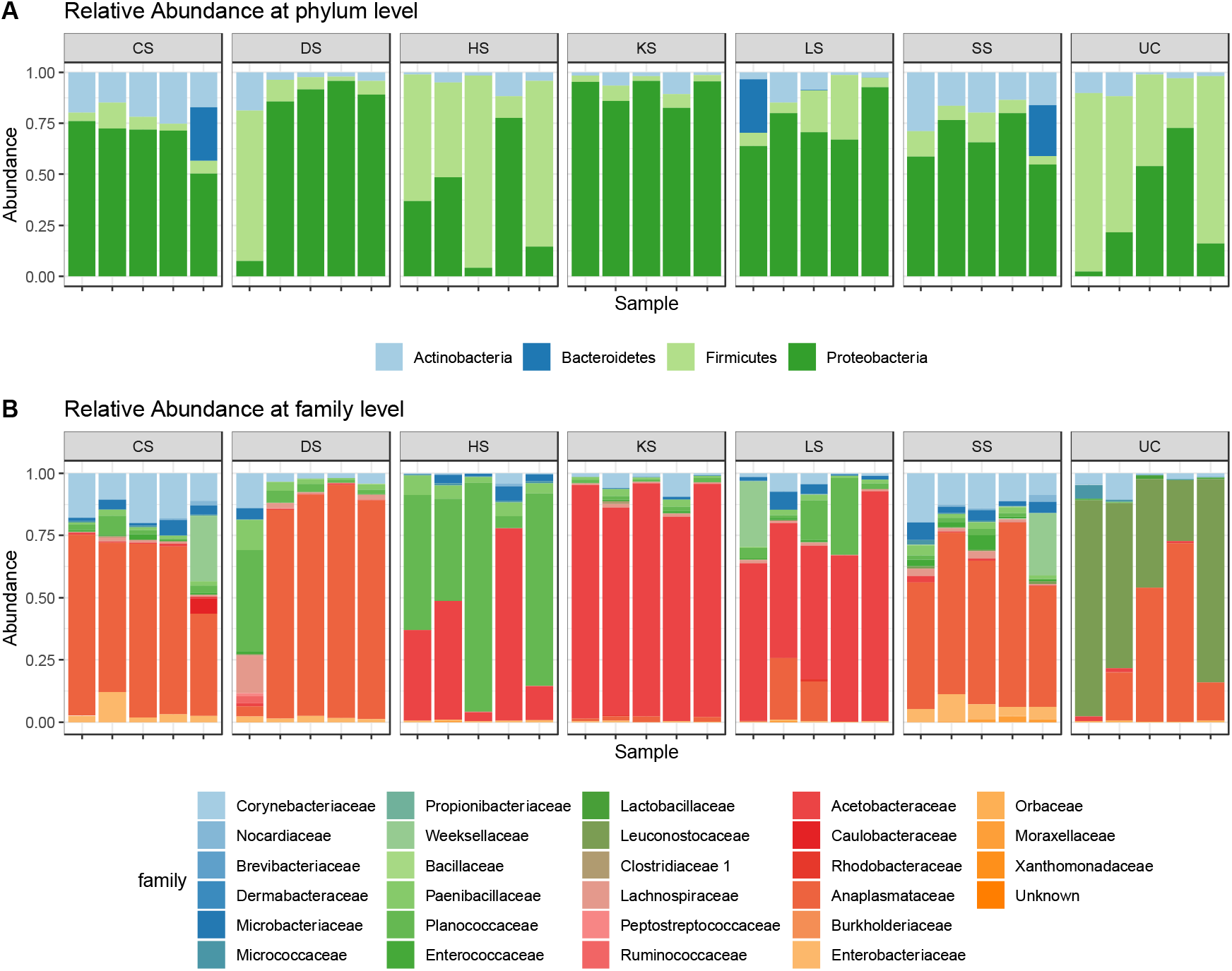
Relative abundance of amplicon sequence variants (ASVs) assigned to the classification level phylum (**A**) and family (**B**) for the six selection regimes (CS: cold-shock resistance, DS: desiccation resistance, HS: heat-shock resistance, KS: heat knockdown resistance, LS: longevity, SS: starvation resistance) and the unselected control lines (UC). Each column within selection type represents a biological replicate.

Comparison across selection lines revealed a change in microbial composition following exposure to artificial selection (Figure 1). At phylum level (Figure 1A), samples separated into two groups – samples characterized by high abundance of Proteobacteria (CS, DS, KS, LS, SS) or high abundance of Firmicutes (HS and UC). At family level (Figure 1B), four microbial profiles were identified: samples with high abundance of i) Anaplasmataceae (CS, DS, SS), ii) Acetobacteraceae (KS, LS), iii) Planococcaceae (HS), or iv) Leuconostocaceae (UC). Moreover, the most abundant genus within each selection line was sufficient to unambiguously separate selection lines into the four microbial profiles detected at family level (Figure S1). Top abundant genera identified included *Wolbachia* representative for the CS, DS, and SS lines, *Acetobacter* representative for KS and LS lines, *Lysinibacillus* representative for the HS line, and *Leuconostoc* representative for the UC line. Compared to the other detected key genera, *Leuconostoc* was exclusively found in the UC lines.

### Inter-sample complexity (α-diversity)

Within-sample diversity was investigated for rarefied and prevalence pruned data, only including ASVs with an overall abundance >0.01% (141 ASVs; range: 9 to 86; mean: 48). Shapiro-Wilk tests confirmed normality of all α-diversity estimates (Observed species richness: *P*=0.80; Shannon’s diversity index: *P*=0.31; Faith’s phylogenetic diversity (PD) index: *P*=0.66, respectively). Global comparison of α-diversity estimates between regimes (six selection regimes and the unselected control regime) showed that α-diversity differed between them (Table 1).

**Table 1.** Analysis of variance for three α-diversity metrics.

|  | Df | Sum Sq | Mean Sq | F value | P value |
| --- | --- | --- | --- | --- | --- |
| **Species richness** |  |  |  |  |  |
| Selection regime | 6 | 9533 | 1588.8 | 12.66 | < 0.001 |
| Residuals | 28 | 3514 | 125.5 |  |  |
| **Shannon's index** |  |  |  |  |  |
| Selection regime | 6 | 6.053 | 1.0089 | 2.775 | < 0.05 |
| Residuals | 28 | 10.179 | 0.3635 |  |  |
| **Faith's Phylogenetic** |  |  |  |  |  |
| Selection regime | 6 | 14.02 | 2.3370 | 13.58 | < 0.001 |
| Residuals | 28 | 4.82 | 0.1721 |  |  |

Df: degrees of freedom; Sq: Squares

Parametric test statistics (Tukey’s HSD test) were applied for pairwise comparison and showed abundant differences in α-diversity metrics between pairs of regimes (Supplementary Tables S1-S3). In general, selection lines showed higher within-sample diversities than the control lines (Figure 2, Supplementary Tables S1-S3).

**Figure 2.**
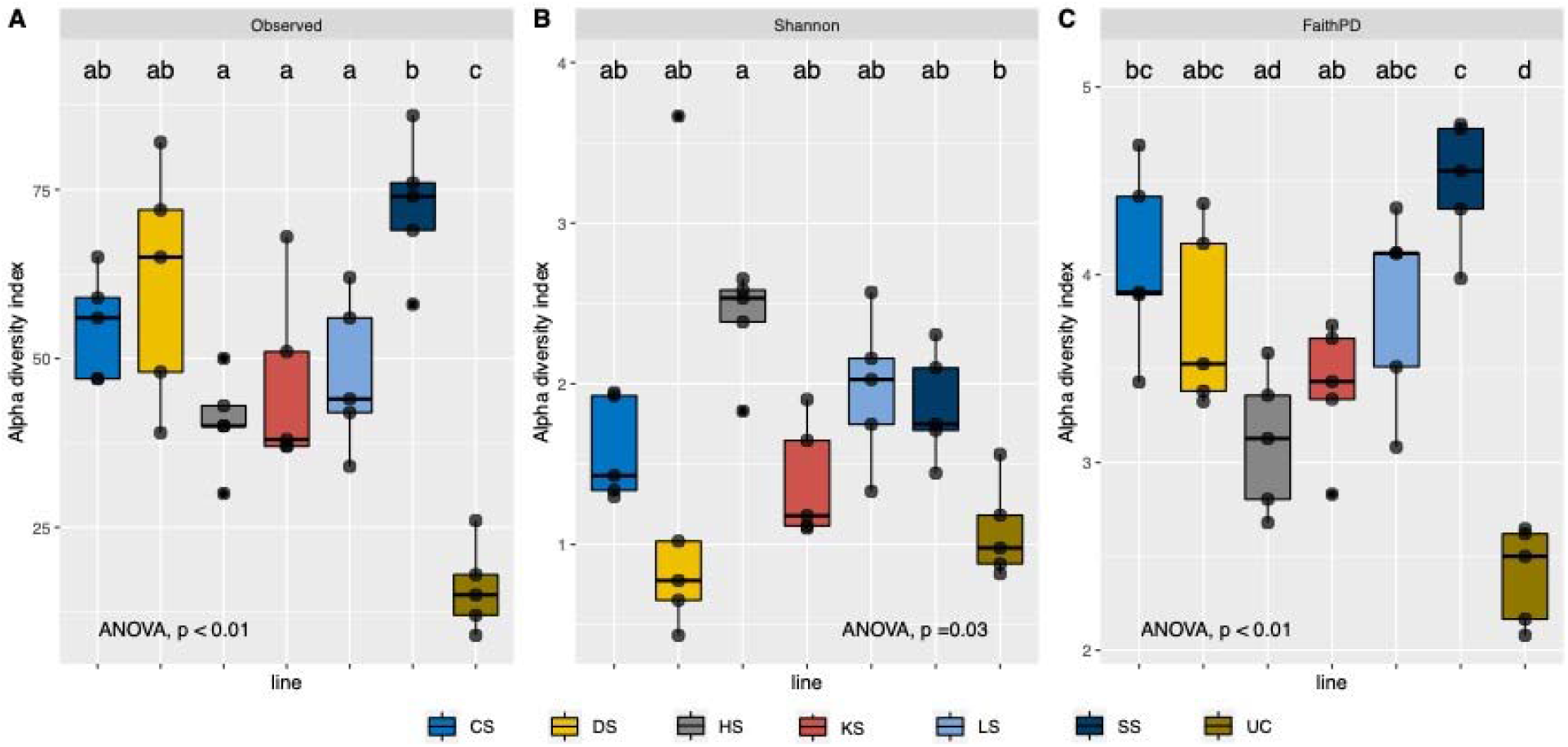
Boxplot for observed (**A**), Shannon’s (**B**) and Faith’s Phylogenetic (**C**) α-diversity metrics. Letters above each boxplot indicate pairwise statistical differences in α-diversity metrics between regimes (CS: cold-shock resistance, DS: desiccation resistance, HS: heat-shock resistance, KS: heat knockdown resistance, LS: longevity, SS: starvation resistance, UC: unselected controls). Full statistical reports are found in Supplementary Tables S1-S3.

Three different α-diversity metrics were estimated to account for differences in richness, evenness, and phylogenetic distance. The observed bacterial richness quantifies the total number of ASV within a sample. The UC line showed significantly less richness compared with the selected lines (Figure 2A). The SS and DS lines displayed highest ASV abundances compared with the other selection lines (Figure 2A). The Shannon’s Index accounts for ASV abundance and evenness, such that samples with comparable numbers of ASVs will show a lower diversity measure when one or few ASVs are overrepresented (uneven distribution). The Shannon’s Index only differed significantly between HS and the control line (Figure 2B), suggesting that ASVs detected in the HS line were more evenly distributed compared to the control line. Finally, the Faith’s phylogenetic diversity metric incorporates the phylogenetic distance between AVSs, such that related ASVs increase the measure of phylogenetic biodiversity less than unrelated ASVs. Like with the observed richness, the control line had a significant lower estimate of Faith’s diversity metric than any of the other selection regimes, and the SS regime displayed the highest diversity metric (Figure 2C).

### Intra-sample complexity (β-diversity)

Among-sample β-diversity was visualized with Principal Coordinate Analysis (PCoA) ordination plots based on Bray-Curtis distances (Figure 3).

**Figure 3.**
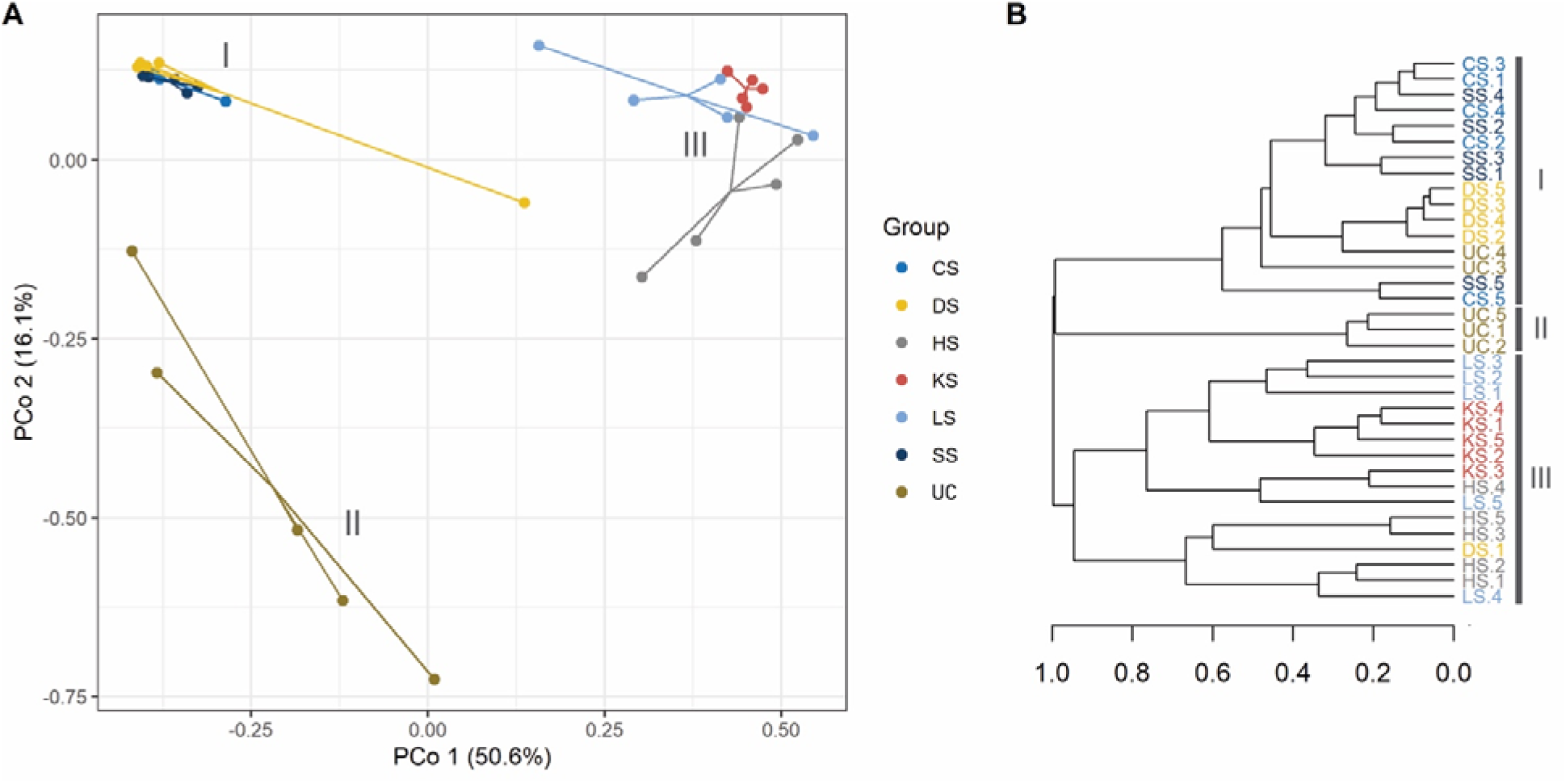
Principal Coordinate (PCo) ordination plot of axis 1 and 2, with proportion of variance explained noted (**A**), and cluster analysis (**B**). Roman numbers indicate distinct separate clusters.

Differences in microbial communities between lines were detected (Figure 3). Based on PCo1 (50.6% of microbial diversity explained) and PCo2 (16.1% of microbial diversity explained), samples separated into three discrete clusters with CS, DS, and SS samples forming cluster I, UC samples forming cluster II, and HS, KS and LS samples forming cluster III. Microbial diversity within the three clusters was highest for UC samples, and lowest among samples belonging to cluster I, indicating a high microbial resemblance among the CS, DS, and SS lines. Permutational multivariate analysis of variance (PERMANOVA) revealed that microbial diversity detected in selection lines clearly differed from that observed in the control lines (Table 3A). Moreover, within-cluster diversity between HS, KS, and LS (cluster III) differed (Table 3C), whereas microbial diversity between CS, DS, and SS (cluster I) did not differ (Table 3B).

**Table 3.** Permutational multivariate analysis of variance (PERMANOVA) for (**A**) selection regimes compared with the unselected control lines, (**B**) pairwise comparison of selection lines belonging to cluster I (CS, DS, SS), and (**C**) pairwise comparison of selection lines belonging to cluster III (HS, KS, LS). PERMANOVA was computed on Bray-Curtis distances.

| **A)** | **Df** | **Sum Sq** | **Mean Sq** | **F value** | **R2** | **Pr(>F)** |
| --- | --- | --- | --- | --- | --- | --- |
| **UC vs. CS** |  |  |  |  |  |  |
| Selection regime | 1 | 0.93624 | 0.93624 | 12.722 | 0.61394 | 0.009 |
| Residuals | 8 | 0.58873 | 0.07359 |  | 0.38606 |  |
| **UC vs. DS** |  |  |  |  |  |  |
| Selection regime | 1 | 0.94849 | 0.94849 | 7.5781 | 0.48646 | 0.013 |
| Residuals | 8 | 1.00129 | 0.12516 |  | 0.51354 |  |
| **UC vs. HS** |  |  |  |  |  |  |
| Selection regime | 1 | 1.95212 | 1.95212 | 16.362 | 0.67162 | 0.012 |
| Residuals | 8 | 0.95447 | 0.11931 |  | 0.32838 |  |
| **UC vs. KS** |  |  |  |  |  |  |
| Selection regime | 1 | 1.99204 | 1.99204 | 24.728 | 0.75556 | 0.004 |
| Residuals | 8 | 0.64446 | 0.08056 |  | 0.24444 |  |
| **UC vs. LS** |  |  |  |  |  |  |
| Selection regime | 10 | 1.69650 | 1.69650 | 14.596 | 0.64596 | 0.01 |
| Residuals | 8 | 0.92983 | 0.11623 |  | 0.35404 |  |
| **UC vs. SS** |  |  |  |  |  |  |
| Selection regime | 1 | 0.95193 | 0.95193 | 14.068 | 0.63748 | 0.012 |
| Residuals | 8 | 0.54134 | 0.06767 |  | 0.36252 |  |
| **B)** | **Df** | **Sum Sq** | **Mean Sq** | **F value** | **R2** | **Pr(>F)** |
| **CS vs. DS** |  |  |  |  |  |  |
| Selection regime | 1 | 0.13416 | 0.134160 | 1.3507 | 0.14445 | 0.22 |
| Residuals | 8 | 0.79458 | 0.099323 |  | 0.85555 |  |
| **CS vs. SS** |  |  |  |  |  |  |
| Selection regime | 1 | 0.03764 | 0.037644 | 0.89994 | 0.10112 | 0.531 |
| Residuals | 8 | 0.33464 | 0.041830 |  | 0.89888 |  |
| **SS vs. DS** |  |  |  |  |  |  |
| Selection regime | 1 | 0.18038 | 0.180377 | 1.9312 | 0.19446 | 0.074 |
| Residuals | 8 | 0.74719 | 0.093399 |  | 0.80554 |  |
| **C)** | **Df** | **Sum Sq** | **Mean Sq** | **F value** | **R2** | **Pr(>F)** |
| **KS vs. LS** |  |  |  |  |  |  |
| Selection regime | 1 | 0.37951 | 0.37951 | 3.8982 | 0.32763 | 0.007 |
| Residuals | 8 | 0.77885 | 0.09736 |  | 0.67237 |  |
| **HS vs. KS** |  |  |  |  |  |  |
| Selection regime | 1 | 0.88727 | 0.88727 | 8.8341 | 0.52478 | 0.009 |
| Residuals | 8 | 0.80350 | 0.10044 |  | 0.47522 |  |
| **HS vs. LS** |  |  |  |  |  |  |
| Selection regime | 1 | 0.55765 | 0.55765 | 4.0971 | 0.33869 | 0.019 |
| Residuals | 8 | 1.08887 | 0.13611 |  | 0.66131 |  |

### Differential abundance

Microbial abundance of the unselected control lines, agglomerated at genus level, was compared with each of the selection regimes, respectively. Of the 32 assigned genera, a total of 20 genera belonging to 13 different families were significantly affected by at least one selection regime (Figure 4).

**Figure 4.**
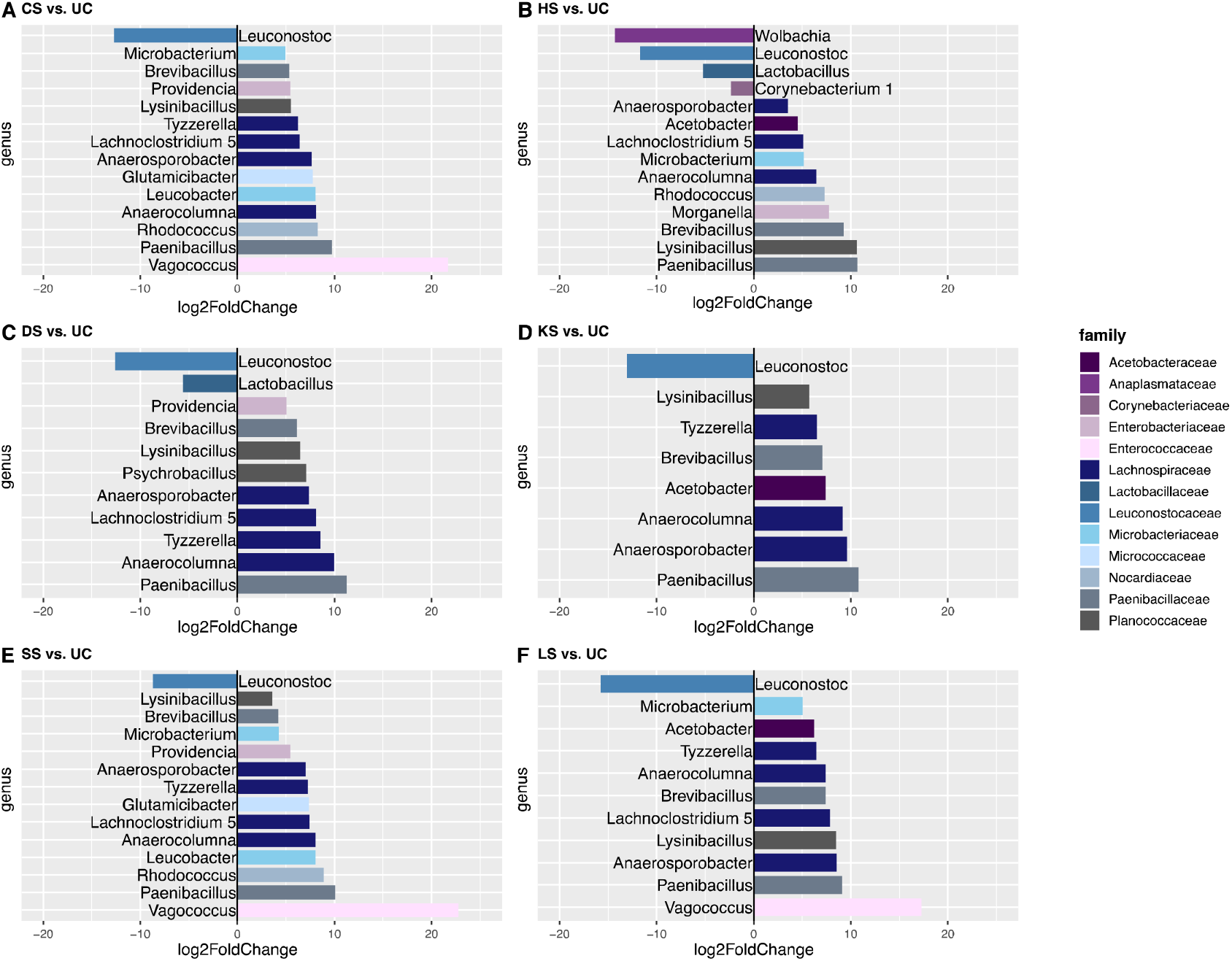
Differential abundance analysis comparing different selection regimes at the genus level. CS: cold-shock resistance, DS: desiccation resistance, HS: heat-shock resistance, KS: heat knockdown resistance, LS: longevity, SS: starvation resistance, UC: unselected controls. Colour code applied represents family assignment. Full statistical reports are found in Supplementary Tables S4-S9.

The number of differentially abundant genera ranged from 8 in the KS line to 14 in the CS, SS, and HS lines. The majority of the differentially detected genera (*Acetobacter*, *Anaerocolumna*, *Anaerosporobacter*, *Brevibacillus*, *Glutamicibacter*, *Lachnoclostridium 5*, *Leucobacter*, *Lysinibacillus*, *Microbacterium*, *Morganella*, *Paenibacillus*, *Providencia*, *Psychrobacillus*, *Rhodococcus*, *Tyzzerella*, *Vagococcus*) showed larger abundances in selection lines, with *Anaerocolumna*, *Anaerosporobacter*, *Brevibacillus*, *Lysinibacillus, and Paenibacillus* being more abundant across all selection regimes, whereas *Morganella* and *Psychrobacillus* were detected differentially abundant in one selection regime (HS; Figure 4). For the remaining four differentially detected genera (*Leuconostoc*, *Lactobacillus*, *Wolbachia* and *Corynebacterium 1*) abundances were lower in selection regimes compared to the control line, with lower *Leuconostoc* abundance detected across all selection regimes, lower *Lactobacillus* abundance detected in the DS and HS selection regimes, and lower *Corynebacterium 1* and *Wolbachia* abundances detected in the HS regime, only. Considering the clusters detected based on β-diversity, *Providencia* was found differentially abundant in all selection regimes belonging to cluster I (CS, DS, SS) but in none of the selection regimes belonging to cluster III (HS, KS, and LS). *Vice versa*, *Acetobacter* was found differentially abundant in all selection regimes belonging to cluster III but in none of the selection regimes belonging to cluster I.

## Discussion

Advances in sequencing technologies have allowed detailed studies of host and microbiome genomes and thereby provided new opportunities for studying interactions between host and associated microorganisms [27,64–67]. Therefore, we can now test the hypothesis that the combination of the host’s and its associated microorganisms’ genomes in concert with environmental variability constitutes an individual’s fitness and comprises the unit of selection. This is the focus of the current study where we utilized a unique set of highly replicated *D. melanogaster* selection and control lines to provide strong experimental support for the hologenome theory of evolution.

We investigated the interactions between host genotype and microbial composition by assessing whether genetic and phenotypic differentiated *D. melanogaster* lines had distinct microbiomes. We investigated microbiomes in replicate control lines, in lines selected for increased longevity, and in lines selected for each of five environmental stress resistance traits, namely increased desiccation resistance (DS), increased cold-shock resistance (CS), increased heat-shock resistance (HS), increased heat knockdown resistance (KS) and increased starvation resistance (SS). Strong functional phenotypic responses to selection in these lines have previously been shown [36,37] along with transcriptome, metabolome, and proteome studies describing the physiological and genomic basis of the response to selection [40–42,44,68]. We found that the microbiome detected in the selection lines differed markedly from that of the control lines (Figures 1–2) and that replicate lines within selection regimes had rather similar microbiomes. This was seen in the form of increased microbial diversity in lines selected for increased longevity or stress resistance and differential abundance of core *D. melanogaster* microbes between selection regimes (Figure 4). We identified three clusters with the CS, SS and DS selection regimes (and two UC lines) constituting cluster I, UC (three of five) lines constituting cluster II and KS, HS, DS and LS selection regimes constituting cluster III (Figure 3). Thus, we convincingly show that the microbial community of the host is strongly linked to fitness components of the host; in our case stress resistance traits and longevity [7–14].

The few previous studies that have investigated the effects of within species host genetic variation on endosymbionts have mainly focused on genetic variability in the host’s ability to control the presence and/or abundance of specific endosymbionts [69,70]. Experimental evidence supporting the hologenome hypothesis, i.e., that the host and microbiome genomes in concert constitute the unit of selection, is however beginning to emerge. For example, Kokou et al. (2018) showed alteration of the microbial composition and microbial response to temperature stress in a tropical fish selected for increased cold resistance, which is in line with the results presented here.

The composition of the host’s microbiome has been linked to variation in the host’s phenotype, such as immune response, metabolism, fitness and developmental characteristics [22,71–73]. The microbiome-associated phenotypic mediations are caused either by the production of metabolites [74] or the presence of the bacteria *per se* [75]. For ectotherms in particular, alteration of energy reserves, metabolism, or gene expression as a consequence of alterations in the microbiome may indirectly affect an individual’s environmental stress resistance [76]. In the current study, we found abundant changes in the microbial community composition between lines under selection and the unselected control lines. For example, we found that Acetobacteraceae were enriched in HS, KS and LS selected regimes (Figure 1 and 4), as predicted from the biochemical mechanisms of Acetobacteraceae, a group of bacteria that reduces lipid storage in the host [73,77]. Consequently, lines with low abundance of Acetobacteraceae have increased lipid storage which is favorable in cold environments [78]. The gram positive Leuconostocaceae, which have been identified in wild-caught *D. melanogaster* [23,79], were underrepresented in all selection lines compared with the unselected control lines (Figure 4). Some members of the *Leuconostoc* genus are known to ferment fructose [80], helping the host with digestion of fruits or other plant materials. Hence, one could speculate whether the costs associated with increased stress resistance [81,82] or longevity [83,84] are associated with reduced ability to process food. *Lactobacillus* has been found to be one of the most common genera within the *Drosophila* microbiome [85], and it has been shown that flies with only *Lactobacillus* microbes can recapitulate the natural microbiota growth-promoting effects [86]. Importantly, chronical stress dramatically changes the microbiota composition and the following metabolomic signature [87]. Here, we found that *Lactobacillus* was less abundant in lines selected for increased heat stress and desiccation resistance (Figure 4). Interestingly, *Wolbachia* was reduced by almost 15-fold in heat stress selected lines compared with unselected control lines (Figure 4). *Wolbachia* is a maternally transmitted bacteria found to infect most insects [88]. Different strains of *Wolbachia* have been shown to have marked effect on temperature preference. *Wolbachia* is generally vulnerable to heat stress [88,89] and certain *Wolbachia* genotypes have been found to promote changes in heat stress resistance of *Drosophila* [90]. During the artificial selection procedures executed to generate the HS lines, flies were exposed to high temperatures that killed a proportion of the population and surviving flies were used to establish the next generation. This heat treatment might have caused high mortality of *Wolbachia*, providing a possible explanation for the lower abundance observed in the HS lines.

Another interesting observation from the HS lines is that we here observed the lowest observed and phylogenetic diversity metrics, but the highest Shannon index (Figure 2). This indicates that only few ASVs were represented and with an uneven distribution. This could be a consequence of the heat exposure selecting for microbes that can tolerate high temperatures (thereby decreasing observed alpha diversity but increasing Shannon’s index). This is supported by our differentially abundance analysis showing that *Paenibacillus* – a heat stable bacteria [91,92] – was the bacteria that was mostly increased in abundance comparing HS and UC (Figure 4).

In our study we reared the 35 *D. melanogaster* lines investigated under common garden conditions and found marked differences in their microbiome. Thus, the fact that certain bacteria are more abundant in some selection regimes suggests that superior host genotypes for e.g., the ability to tolerate low temperatures (CS lines) provide competitive advantages of different microbes compared to those in the non-selected control lines. Our data suggests that as the host evolves so does the microbiome and that it is the combined effect of host and microbial evolution that governs the functional phenotypic responses to artificial selection observed in the investigated lines [36,37]. Verifying this hypothesis obviously demands further investigations including microbiome transfer and manipulation experiments along the lines proposed by Moghadam et al. [13] and Ericsson and Franklin [9]. In interpreting our results it is also important to take into account that the relationship between resident microbes and host physiology is very complex and is being influenced by a range of factors including host genotype [93], immunity [94], diet [95] and interactions between microbes [96].

## Acknowledgements

This work was supported by the Danish Council for Independent Research (TNK: grant #DFF-8021-00014b, VL: grant #DFF-4002-00113b).

## Author contributions

T.N.K. and V.L. conceived the ideas and designed the methodology. J.G.S. and V.L. contributed to establishment of the lines. A.S. and P.D.R. analyzed data. T.N.K., A.S. and P.D.R. led the writing of the manuscript and all authors contributed to the drafts and gave final approval for publication.

