## Supplementary Material for "Strong experimental support for the hologenome hypothesis revealed from *Drosophila melanogaster* selection lines"

*Shared first authorship

† Corresponding author

The supplementary material contains the following:

**Supplementary Figures**

Figure S1.	Relative abundance (%) of amplicon sequence variants (ASVs) assigned to family level.

**Supplementary Tables**

Table S1.	Parametric pairwise test statistics for observed diversity.

Table S2.	Parametric pairwise test statistics for Shannon’s diversity index.

Table S3.	Parametric pairwise test statistics for Faith´s phylogenetic diversity index.

Table S4.	Differential abundance analysis comparing

Table S5.	Differential abundance analysis comparing lines selected for desiccation resistance (DS) to the unselected control lines (UC) at the genus level.

Table S6.	Differential abundance analysis comparing lines selected for heat-shock resistance (HS) to the unselected control lines (UC) at the genus level.

Table S7.	Differential abundance analysis comparing lines selected for heat knockdown resistance (KS) to the unselected control lines (UC) at the genus level.

Table S8.	Differential abundance analysis comparing lines selected for longevity (LS) to the unselected control lines (UC) at the genus level.

Table S9.	Differential abundance analysis comparing lines selected for starvation resistance (SS) to the unselected control lines (UC) at the genus level.

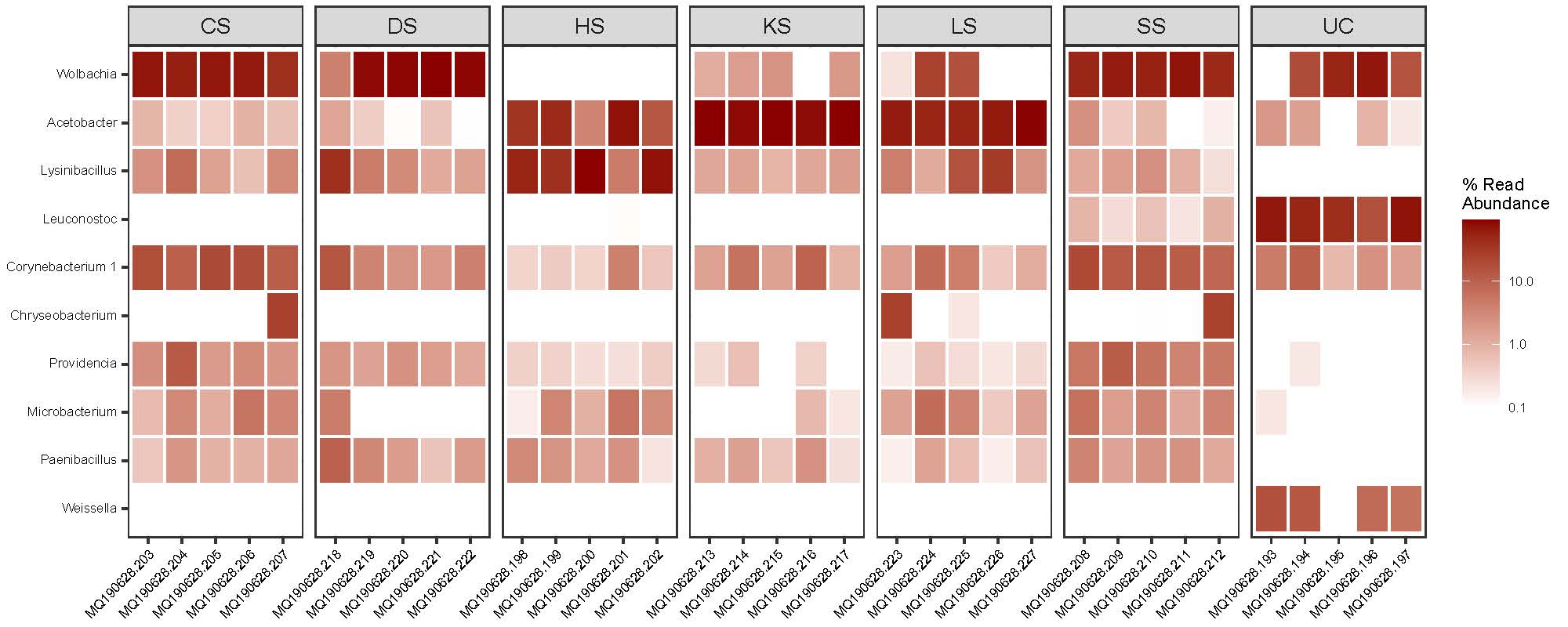

**Figure S1**. Relative abundance (%) of amplicon sequence variants (ASVs) assigned to family level for the six selection regimes (CS: cold-shock resistance, DS: desiccation resistance, HS: heat-shock resistance, KS: heat knockdown resistance, LS: longevity, SS: starvation resistance) and the unselected control lines (UC). Each column within selection type represents the one of the five biological replicates.

**Table S1**. Parametric pairwise test statistics (Tukey´s HSD) for observed $\alpha$-diversity. Rows in bold indicate statistically significant differences between pairs of selection regimes.

| **Comparison** | **Δα** | **95% confidence interval** | | ***P* adj** |
| --- | --- | --- | --- | --- |
| DS-CS | 6.4 | -16.075 | 28.875 | 0.969 |
| HS-CS | -14.20 | -36.675 | 8.275 | 0.434 |
| KS-CS | -8.60 | -31.075 | 13.875 | 0.883 |
| LS-CS | -7.20 | -29.675 | 15.275 | 0.946 |
| SS-CS | 17.80 | -4.675 | 40.275 | 0.193 |
| **UC-CS** | **-38.80** | **-61.275** | **-16.325** | **1.40E-04** |
| HS-DS | -20.60 | -43.075 | 1.875 | 0.089 |
| KS-DS | -15.00 | -37.475 | 7.475 | 0.371 |
| LS-DS | -13.60 | -36.075 | 8.875 | 0.485 |
| SS-DS | 11.40 | -11.075 | 33.875 | 0.678 |
| **UC-DS** | **-45.20** | **-67.675** | **-22.725** | **1.27E-05** |
| KS-HS | 5.60 | -16.875 | 28.075 | 0.984 |
| LS-HS | 7.00 | -15.475 | 29.475 | 0.953 |
| **SS-HS** | **32.00** | **9.525** | **54.475** | **0.002** |
| **UC-HS** | **-24.60** | **-47.075** | **-2.125** | **0.025** |
| LS-KS | 1.40 | -21.075 | 23.875 | 1.000 |
| **SS-KS** | 26.40 | 3.925 | 48.875 | 0.014 |
| **UC-KS** | **-30.20** | **-52.675** | **-7.725** | **0.003** |
| SS-LS | 25.00 | 2.525 | 47.475 | 0.022 |
| **UC-LS** | **-31.60** | **-54.075** | **-9.125** | **0.002** |
| **UC-SS** | **-56.60** | **-79.075** | **-34.125** | **2.10E-07** |
| $\Delta\alpha$ difference $\alpha$-diversity between pairs of selection regimes. | | | | |

**Table S2**. Parametric pairwise test statistics (Tukey´s HSD) for Shannon’s $\alpha$-diversity index. Rows in bold indicate statistically significant differences between pairs of selection regimes.

| **Comparison** | **Δα** | **95% confidence interval** | | ***P* adj** |
| --- | --- | --- | --- | --- |
| DS-CS | -0.28 | -1.49 | 0.93 | 0.99 |
| HS-CS | 0.81 | -0.40 | 2.02 | 0.37 |
| KS-CS | -0.20 | -1.41 | 1.01 | 1.00 |
| LS-CS | 0.38 | -0.83 | 1.59 | 0.95 |
| SS-CS | 0.27 | -0.94 | 1.48 | 0.99 |
| UC-CS | -0.50 | -1.71 | 0.71 | 0.84 |
| HS-DS | 1.09 | -0.12 | 2.30 | 0.10 |
| KS-DS | 0.08 | -1.13 | 1.29 | 1.00 |
| LS-DS | 0.66 | -0.55 | 1.87 | 0.61 |
| SS-DS | 0.55 | -0.66 | 1.76 | 0.77 |
| UC-DS | -0.23 | -1.44 | 0.98 | 1.00 |
| KS-HS | -1.01 | -2.22 | 0.20 | 0.15 |
| LS-HS | -0.43 | -1.64 | 0.78 | 0.91 |
| SS-HS | -0.54 | -1.74 | 0.67 | 0.80 |
| **UC-HS** | **-1.31** | **-2.52** | **-0.10** | **0.03** |
| LS-KS | 0.58 | -0.63 | 1.79 | 0.73 |
| SS-KS | 0.47 | -0.74 | 1.68 | 0.87 |
| UC-KS | -0.31 | -1.51 | 0.90 | 0.98 |
| SS-LS | -0.10 | -1.31 | 1.10 | 1.00 |
| UC-LS | -0.88 | -2.09 | 0.33 | 0.27 |
| UC-SS | -0.78 | -1.99 | 0.43 | 0.41 |
| $\Delta\alpha$ difference $\alpha$-diversity between pairs of selection regimes. | | | | |

**Table S3**. Parametric pairwise test statistics (Tukey´s HSD) for Faith´s phylogenetic diversity. Rows in bold indicate statistically significant differences between pairs of selection regimes.

| **Comparison** | **Δα** | **95% confidence interval** | | ***P* adj** |
| --- | --- | --- | --- | --- |
| DS-CS | -0.31 | -1.14 | 0.52 | 0.89 |
| HS-CS | -0.96 | -1.79 | -0.12 | 0.02 |
| KS-CS | -0.67 | -1.50 | 0.16 | 0.18 |
| LS-CS | -0.23 | -1.06 | 0.60 | 0.97 |
| SS-CS | 0.43 | -0.41 | 1.26 | 0.67 |
| **UC-CS** | **-1.66** | **-2.50** | **-0.83** | **1.40E-05** |
| HS-DS | -0.64 | -1.48 | 0.19 | 0.21 |
| KS-DS | -0.36 | -1.19 | 0.48 | 0.82 |
| LS-DS | 0.08 | -0.75 | 0.91 | 1.00 |
| SS-DS | 0.74 | -0.10 | 1.57 | 0.11 |
| **UC-DS** | **-1.35** | **-2.18** | **-0.52** | **3.30E-04** |
| KS-HS | 0.29 | -0.54 | 1.12 | 0.92 |
| LS-HS | 0.72 | -0.11 | 1.56 | 0.12 |
| **SS-HS** | **1.38** | **0.55** | **2.21** | **2.45E-04** |
| UC-HS | -0.71 | -1.54 | 0.12 | 0.14 |
| LS-KS | 0.44 | -0.40 | 1.27 | 0.64 |
| **SS-KS** | **1.09** | **0.26** | **1.93** | **4.43E-03** |
| UC-KS | -1.00 | -1.83 | -0.16 | 0.01 |
| SS-LS | 0.66 | -0.18 | 1.49 | 0.20 |
| **UC-LS** | **-1.43** | **-2.26** | **-0.60** | **1.46E-04** |
| **UC-SS** | **-2.09** | **-2.92** | **-1.26** | **2.24E-07** |
| $\Delta\alpha$ difference $\alpha$-diversity between pairs of selection regimes. | | | | |

**Table S4**. Differential abundance analysis comparing lines selected for cold-shock resistance (CS) to the unselected control lines (UC) at the genus level. Taxonomies were agglomerated to genus level prior to analysis. ASVs are ordered according to adjusted P-values (P adj) from lowest to highest. ASVs were considered differentially abundant for adjusted P-values ≤ 0.01 and an FDR cut-off of 5%. Significance threshold (P adj = 0.01) is marked by a dashed line.

| **ASV ID** | **baseMean** | **log2FC** | ***P* adj** | **Domain** | **Phylum** | **Class** | **Order** | **Family** | **Genus** |
| --- | --- | --- | --- | --- | --- | --- | --- | --- | --- |
| abc8f555ea0b005c9a4085a8343a3f99 | 159.83 | 9.71 | 9.58E-23 | Bacteria | Firmicutes | Bacilli | Bacillales | Paenibacillaceae | Paenibacillus |
| 810bf4f2547219682f5ab14a09ca3599 | 2656.56 | -12.72 | 7.81E-18 | Bacteria | Firmicutes | Bacilli | Lactobacillales | Leuconostocaceae | Leuconostoc |
| 33b3b41ddb67bb49f482abc6126da323 | 32.73 | 7.63 | 1.05E-12 | Bacteria | Firmicutes | Clostridia | Clostridiales | Lachnospiraceae | Anaerosporobacter |
| ebc7527f35924613e64e071623af1abb | 53.37 | 8.10 | 8.81E-09 | Bacteria | Firmicutes | Clostridia | Clostridiales | Lachnospiraceae | Anaerocolumna |
| 19492e6d689a95489805fe62a81b5ecd | 108.06 | 5.32 | 6.47E-08 | Bacteria | Firmicutes | Bacilli | Bacillales | Paenibacillaceae | Brevibacillus |
| c37413af5ea0a7254284c92b8bf54ae7 | 225.43 | 5.46 | 2.17E-07 | Bacteria | Proteobacteria | Gammaproteobacteria | Enterobacteriales | Enterobacteriaceae | Providencia |
| 05f5c262eb96bf78b6ab2750fdddc0c4 | 18.05 | 6.40 | 5.92E-06 | Bacteria | Firmicutes | Clostridia | Clostridiales | Lachnospiraceae | Lachnoclostridium 5 |
| 5634c39a6b4a1475938b6ad7a4113445 | 14.10 | 21.72 | 9.89E-06 | Bacteria | Firmicutes | Bacilli | Lactobacillales | Enterococcaceae | Vagococcus |
| 148a7bc57792072043058510530ea32d | 2331.90 | 5.53 | 1.68E-05 | Bacteria | Firmicutes | Bacilli | Bacillales | Planococcaceae | Lysinibacillus |
| 7406188bce39180c4aa194bfb387935e | 18.48 | 6.22 | 9.17E-05 | Bacteria | Firmicutes | Clostridia | Clostridiales | Lachnospiraceae | Tyzzerella |
| 9eadd1514e5fcfa10e723634e35f3485 | 197.83 | 4.96 | 5.86E-04 | Bacteria | Actinobacteria | Actinobacteria | Micrococcales | Microbacteriaceae | Microbacterium |
| c2b73b7534124edd282fa67a545e048b | 25.89 | 8.26 | 8.42E-04 | Bacteria | Actinobacteria | Actinobacteria | Corynebacteriales | Nocardiaceae | Rhodococcus |
| c63d552898d6b61e6ef07797210d7aca | 14.67 | 8.05 | 1.66E-03 | Bacteria | Actinobacteria | Actinobacteria | Micrococcales | Microbacteriaceae | Leucobacter |
| cd416ce5bf4afa614154fc0a45feb5a8 | 12.51 | 7.77 | 1.83E-03 | Bacteria | Actinobacteria | Actinobacteria | Micrococcales | Micrococcaceae | Glutamicibacter |
| 64bfc35722d08fe4ccf6b5665b9d1769 | 7.70 | 5.99 | 1.59E-02 | Bacteria | Proteobacteria | Gammaproteobacteria | Enterobacteriales | Enterobacteriaceae | Morganella |
| 19019fa034a9856e8d6365d34574c8fd | 324.39 | -13.68 | 2.18E-02 | Bacteria | Firmicutes | Bacilli | Lactobacillales | Leuconostocaceae | Weissella |
| ffaedab25be5e2be043b2355be6644cf | 9.83 | 5.84 | 6.34E-02 | Bacteria | Firmicutes | Clostridia | Clostridiales | Clostridiaceae 1 | Clostridium sensu stricto 1 |
| 4e817d860267844d0bb67bb3ed49522c | 16.36 | -2.88 | 9.57E-02 | Bacteria | Firmicutes | Bacilli | Lactobacillales | Lactobacillaceae | Lactobacillus |
| 7fda73c8ad009ec246fc6a5bf669203e | 860.52 | 1.43 | 9.79E-02 | Bacteria | Actinobacteria | Actinobacteria | Corynebacteriales | Corynebacteriaceae | Corynebacterium 1 |
| f8d46ee1054ee9224a7e622fd2557cf4 | 3.91 | 4.59 | 1.83E-01 | Bacteria | Firmicutes | Bacilli | Bacillales | Paenibacillaceae | Cohnella |
| f580b2c49c0078358680733c5d838ca6 | 7.02 | 3.55 | 1.98E-01 | Bacteria | Firmicutes | Bacilli | Bacillales | Planococcaceae | Psychrobacillus |
| 0ff98df849b8fae49800a796dadc506c | 49.70 | 1.04 | 3.63E-01 | Bacteria | Firmicutes | Bacilli | Lactobacillales | Enterococcaceae | Enterococcus |
| f8a687d17d5bcd547799c35ad359bf70 | 6790.79 | -0.88 | 3.81E-01 | Bacteria | Proteobacteria | Alphaproteobacteria | Acetobacterales | Acetobacteraceae | Acetobacter |
| a406a791970c2c9105ced42674853b40 | 5366.05 | 1.25 | 4.34E-01 | Bacteria | Proteobacteria | Alphaproteobacteria | Rickettsiales | Anaplasmataceae | Wolbachia |
| b9a736421dc15f263c93ec9f509d5f6b | 2.97 | 2.39 | 7.40E-01 | Bacteria | Firmicutes | Clostridia | Clostridiales | Peptostreptococcaceae | Clostridioides |
| 43f9d96deeccbd4b7c28e80612ec05ea | 4.96 | -1.01 | 7.58E-01 | Bacteria | Actinobacteria | Actinobacteria | Micrococcales | Brevibacteriaceae | Brevibacterium |
| b5e95cda7f3db71495303d91e0c9dda7 | 11.99 | 0.97 | 8.60E-01 | Bacteria | Proteobacteria | Gammaproteobacteria | Pseudomonadales | Moraxellaceae | Acinetobacter |
| 515ecf421a951bc274a7daaa791dc894 | 2.76 | 0.00 | 1.00E+00 | Bacteria | Proteobacteria | Alphaproteobacteria | Rhodobacterales | Rhodobacteraceae | Paracoccus |

baseMean: Mean abundance of the normalized count values over all samples included in the comparison; log2FC: log2-fold change represents the effect size estimate reported on a logarithmic scale to base 2, indicating how much the ASV abundance has changed due to selection for cold-shock resistance in comparison to the control lines; *P* adj: adjusted P-values using the Benjamini-Hochberg procedure.

**Table S5**. Differential abundance analysis comparing lines selected for desiccation resistance (DS) to the unselected control lines (UC) at the genus level. Taxonomies were agglomerated to genus level prior to analysis. ASVs are ordered according to adjusted P-values (P adj) from lowest to highest. ASVs were considered differentially abundant for adjusted P-values ≤ 0.01 and an FDR cut-off of 5%. Significance threshold (P adj = 0.01) is marked by a dashed line.

| **ASV ID** | **baseMean** | **log2FC** | ***P* adj** | **Domain** | **Phylum** | **Class** | **Order** | **Family** | **Genus** |
| --- | --- | --- | --- | --- | --- | --- | --- | --- | --- |
| abc8f555ea0b005c9a4085a8343a3f99 | 159.83 | 11.23 | 2.10E-30 | Bacteria | Firmicutes | Bacilli | Bacillales | Paenibacillaceae | Paenibacillus |
| 810bf4f2547219682f5ab14a09ca3599 | 2656.56 | -12.60 | 4.06E-17 | Bacteria | Firmicutes | Bacilli | Lactobacillales | Leuconostocaceae | Leuconostoc |
| ebc7527f35924613e64e071623af1abb | 53.37 | 9.95 | 7.92E-13 | Bacteria | Firmicutes | Clostridia | Clostridiales | Lachnospiraceae | Anaerocolumna |
| 33b3b41ddb67bb49f482abc6126da323 | 32.73 | 7.34 | 8.62E-12 | Bacteria | Firmicutes | Clostridia | Clostridiales | Lachnospiraceae | Anaerosporobacter |
| 19492e6d689a95489805fe62a81b5ecd | 108.06 | 6.12 | 3.16E-10 | Bacteria | Firmicutes | Bacilli | Bacillales | Paenibacillaceae | Brevibacillus |
| 05f5c262eb96bf78b6ab2750fdddc0c4 | 18.05 | 8.08 | 5.13E-09 | Bacteria | Firmicutes | Clostridia | Clostridiales | Lachnospiraceae | Lachnoclostridium 5 |
| 7406188bce39180c4aa194bfb387935e | 18.48 | 8.54 | 4.22E-08 | Bacteria | Firmicutes | Clostridia | Clostridiales | Lachnospiraceae | Tyzzerella |
| 148a7bc57792072043058510530ea32d | 2331.90 | 6.48 | 3.66E-07 | Bacteria | Firmicutes | Bacilli | Bacillales | Planococcaceae | Lysinibacillus |
| c37413af5ea0a7254284c92b8bf54ae7 | 225.43 | 5.05 | 1.40E-06 | Bacteria | Proteobacteria | Gammaproteobacteria | Enterobacteriales | Enterobacteriaceae | Providencia |
| 4e817d860267844d0bb67bb3ed49522c | 16.36 | -5.61 | 1.88E-03 | Bacteria | Firmicutes | Bacilli | Lactobacillales | Lactobacillaceae | Lactobacillus |
| f580b2c49c0078358680733c5d838ca6 | 7.02 | 7.09 | 9.36E-03 | Bacteria | Firmicutes | Bacilli | Bacillales | Planococcaceae | Psychrobacillus |
| 19019fa034a9856e8d6365d34574c8fd | 324.39 | -13.31 | 3.74E-02 | Bacteria | Firmicutes | Bacilli | Lactobacillales | Leuconostocaceae | Weissella |
| ffaedab25be5e2be043b2355be6644cf | 9.83 | 4.97 | 1.83E-01 | Bacteria | Firmicutes | Clostridia | Clostridiales | Clostridiaceae 1 | Clostridium sensu stricto 1 |
| a406a791970c2c9105ced42674853b40 | 5366.05 | 2.27 | 2.26E-01 | Bacteria | Proteobacteria | Alphaproteobacteria | Rickettsiales | Anaplasmataceae | Wolbachia |
| 9eadd1514e5fcfa10e723634e35f3485 | 197.83 | 1.79 | 3.49E-01 | Bacteria | Actinobacteria | Actinobacteria | Micrococcales | Microbacteriaceae | Microbacterium |
| f8a687d17d5bcd547799c35ad359bf70 | 6790.79 | -1.16 | 3.49E-01 | Bacteria | Proteobacteria | Alphaproteobacteria | Acetobacterales | Acetobacteraceae | Acetobacter |
| b9a736421dc15f263c93ec9f509d5f6b | 2.97 | 6.22 | 4.18E-01 | Bacteria | Firmicutes | Clostridia | Clostridiales | Peptostreptococcaceae | Clostridioides |
| 43f9d96deeccbd4b7c28e80612ec05ea | 4.96 | -3.21 | 4.18E-01 | Bacteria | Actinobacteria | Actinobacteria | Micrococcales | Brevibacteriaceae | Brevibacterium |
| 016b118f0377208f78e5694abe0d59b6 | 2.09 | 6.02 | 4.27E-01 | Bacteria | Firmicutes | Clostridia | Clostridiales | Ruminococcaceae | Ruminiclostridium |
| 69052d7a94480e1c87649a5aaf431114 | 1.21 | 2.80 | 8.07E-01 | Bacteria | Firmicutes | Clostridia | Clostridiales | Lachnospiraceae | Lachnospiraceae NC2004 group |
| 7fda73c8ad009ec246fc6a5bf669203e | 860.52 | -0.45 | 8.08E-01 | Bacteria | Actinobacteria | Actinobacteria | Corynebacteriales | Corynebacteriaceae | Corynebacterium 1 |
| 0ff98df849b8fae49800a796dadc506c | 49.70 | -0.50 | 8.36E-01 | Bacteria | Firmicutes | Bacilli | Lactobacillales | Enterococcaceae | Enterococcus |
| c2b73b7534124edd282fa67a545e048b | 25.89 | 0.00 | 1.00E+00 | Bacteria | Actinobacteria | Actinobacteria | Corynebacteriales | Nocardiaceae | Rhodococcus |
| cd416ce5bf4afa614154fc0a45feb5a8 | 12.51 | 0.00 | 1.00E+00 | Bacteria | Actinobacteria | Actinobacteria | Micrococcales | Micrococcaceae | Glutamicibacter |
| c63d552898d6b61e6ef07797210d7aca | 14.67 | 0.00 | 1.00E+00 | Bacteria | Actinobacteria | Actinobacteria | Micrococcales | Microbacteriaceae | Leucobacter |
| b5e95cda7f3db71495303d91e0c9dda7 | 11.99 | 0.00 | 1.00E+00 | Bacteria | Proteobacteria | Gammaproteobacteria | Pseudomonadales | Moraxellaceae | Acinetobacter |
| 64bfc35722d08fe4ccf6b5665b9d1769 | 7.70 | 0.00 | 1.00E+00 | Bacteria | Proteobacteria | Gammaproteobacteria | Enterobacteriales | Enterobacteriaceae | Morganella |
| 515ecf421a951bc274a7daaa791dc894 | 2.76 | 0.00 | 1.00E+00 | Bacteria | Proteobacteria | Alphaproteobacteria | Rhodobacterales | Rhodobacteraceae | Paracoccus |
| f8d46ee1054ee9224a7e622fd2557cf4 | 3.91 | 0.00 | 1.00E+00 | Bacteria | Firmicutes | Bacilli | Bacillales | Paenibacillaceae | Cohnella |
| 5634c39a6b4a1475938b6ad7a4113445 | 14.10 | 0.00 | 1.00E+00 | Bacteria | Firmicutes | Bacilli | Lactobacillales | Enterococcaceae | Vagococcus |

baseMean: Mean abundance of the normalized count values over all samples included in the comparison; log2FC: log2-fold change represents the effect size estimate reported on a logarithmic scale to base 2, indicating how much the ASV abundance has changed due to selection for desiccation resistance in comparison to the control lines; *P* adj: adjusted P-values using the Benjamini-Hochberg procedure.

**Table S6**. Differential abundance analysis comparing lines selected for heat-shock resistance (HS) to the unselected control lines (UC) at the genus level. Taxonomies were agglomerated to genus level prior to analysis. ASVs are ordered according to adjusted P-values (P adj) from lowest to highest. ASVs were considered differentially abundant for adjusted P-values ≤ 0.01 and an FDR cut-off of 5%. Significance threshold (P adj = 0.01) is marked by a dashed line.

| **ASV ID** | **baseMean** | **log2FC** | ***P* adj** | **Domain** | **Phylum** | **Class** | **Order** | **Family** | **Genus** |
| --- | --- | --- | --- | --- | --- | --- | --- | --- | --- |
| abc8f555ea0b005c9a4085a8343a3f99 | 159.83 | 10.66 | 2.18E-27 | Bacteria | Firmicutes | Bacilli | Bacillales | Paenibacillaceae | Paenibacillus |
| 19492e6d689a95489805fe62a81b5ecd | 108.06 | 9.30 | 1.44E-22 | Bacteria | Firmicutes | Bacilli | Bacillales | Paenibacillaceae | Brevibacillus |
| 148a7bc57792072043058510530ea32d | 2331.90 | 10.60 | 2.52E-17 | Bacteria | Firmicutes | Bacilli | Bacillales | Planococcaceae | Lysinibacillus |
| a406a791970c2c9105ced42674853b40 | 5366.05 | -14.29 | 6.67E-17 | Bacteria | Proteobacteria | Alphaproteobacteria | Rickettsiales | Anaplasmataceae | Wolbachia |
| 810bf4f2547219682f5ab14a09ca3599 | 2656.56 | -11.68 | 6.55E-16 | Bacteria | Firmicutes | Bacilli | Lactobacillales | Leuconostocaceae | Leuconostoc |
| f8a687d17d5bcd547799c35ad359bf70 | 6790.79 | 4.55 | 8.46E-07 | Bacteria | Proteobacteria | Alphaproteobacteria | Acetobacterales | Acetobacteraceae | Acetobacter |
| ebc7527f35924613e64e071623af1abb | 53.37 | 6.46 | 6.71E-06 | Bacteria | Firmicutes | Clostridia | Clostridiales | Lachnospiraceae | Anaerocolumna |
| 9eadd1514e5fcfa10e723634e35f3485 | 197.83 | 5.16 | 4.86E-04 | Bacteria | Actinobacteria | Actinobacteria | Micrococcales | Microbacteriaceae | Microbacterium |
| 05f5c262eb96bf78b6ab2750fdddc0c4 | 18.05 | 5.12 | 5.14E-04 | Bacteria | Firmicutes | Clostridia | Clostridiales | Lachnospiraceae | Lachnoclostridium 5 |
| 64bfc35722d08fe4ccf6b5665b9d1769 | 7.70 | 7.75 | 1.97E-03 | Bacteria | Proteobacteria | Gammaproteobacteria | Enterobacteriales | Enterobacteriaceae | Morganella |
| 4e817d860267844d0bb67bb3ed49522c | 16.36 | -5.21 | 3.21E-03 | Bacteria | Firmicutes | Bacilli | Lactobacillales | Lactobacillaceae | Lactobacillus |
| c2b73b7534124edd282fa67a545e048b | 25.89 | 7.30 | 4.09E-03 | Bacteria | Actinobacteria | Actinobacteria | Corynebacteriales | Nocardiaceae | Rhodococcus |
| 33b3b41ddb67bb49f482abc6126da323 | 32.73 | 3.52 | 4.70E-03 | Bacteria | Firmicutes | Clostridia | Clostridiales | Lachnospiraceae | Anaerosporobacter |
| 7fda73c8ad009ec246fc6a5bf669203e | 860.52 | -2.33 | 6.23E-03 | Bacteria | Actinobacteria | Actinobacteria | Corynebacteriales | Corynebacteriaceae | Corynebacterium 1 |
| 0ff98df849b8fae49800a796dadc506c | 49.70 | -2.70 | 1.57E-02 | Bacteria | Firmicutes | Bacilli | Lactobacillales | Enterococcaceae | Enterococcus |
| 19019fa034a9856e8d6365d34574c8fd | 324.39 | -13.51 | 2.55E-02 | Bacteria | Firmicutes | Bacilli | Lactobacillales | Leuconostocaceae | Weissella |
| c37413af5ea0a7254284c92b8bf54ae7 | 225.43 | 2.28 | 4.04E-02 | Bacteria | Proteobacteria | Gammaproteobacteria | Enterobacteriales | Enterobacteriaceae | Providencia |
| cd416ce5bf4afa614154fc0a45feb5a8 | 12.51 | 4.90 | 6.02E-02 | Bacteria | Actinobacteria | Actinobacteria | Micrococcales | Micrococcaceae | Glutamicibacter |
| f8d46ee1054ee9224a7e622fd2557cf4 | 3.91 | 6.31 | 6.02E-02 | Bacteria | Firmicutes | Bacilli | Bacillales | Paenibacillaceae | Cohnella |
| ffaedab25be5e2be043b2355be6644cf | 9.83 | 4.58 | 1.59E-01 | Bacteria | Firmicutes | Clostridia | Clostridiales | Clostridiaceae 1 | Clostridium sensu stricto 1 |
| 7406188bce39180c4aa194bfb387935e | 18.48 | 2.50 | 1.79E-01 | Bacteria | Firmicutes | Clostridia | Clostridiales | Lachnospiraceae | Tyzzerella |
| b5e95cda7f3db71495303d91e0c9dda7 | 11.99 | 4.27 | 4.58E-01 | Bacteria | Proteobacteria | Gammaproteobacteria | Pseudomonadales | Moraxellaceae | Acinetobacter |
| f580b2c49c0078358680733c5d838ca6 | 7.02 | 0.00 | 1.00E+00 | Bacteria | Firmicutes | Bacilli | Bacillales | Planococcaceae | Psychrobacillus |
| 69052d7a94480e1c87649a5aaf431114 | 1.21 | 0.00 | 1.00E+00 | Bacteria | Firmicutes | Clostridia | Clostridiales | Lachnospiraceae | Lachnospiraceae NC2004 group |
| b9a736421dc15f263c93ec9f509d5f6b | 2.97 | 0.00 | 1.00E+00 | Bacteria | Firmicutes | Clostridia | Clostridiales | Peptostreptococcaceae | Clostridioides |
| 016b118f0377208f78e5694abe0d59b6 | 2.09 | 0.00 | 1.00E+00 | Bacteria | Firmicutes | Clostridia | Clostridiales | Ruminococcaceae | Ruminiclostridium |
| c63d552898d6b61e6ef07797210d7aca | 14.67 | 0.00 | 1.00E+00 | Bacteria | Actinobacteria | Actinobacteria | Micrococcales | Microbacteriaceae | Leucobacter |
| 43f9d96deeccbd4b7c28e80612ec05ea | 4.96 | 0.59 | 1.00E+00 | Bacteria | Actinobacteria | Actinobacteria | Micrococcales | Brevibacteriaceae | Brevibacterium |
| 515ecf421a951bc274a7daaa791dc894 | 2.76 | 0.00 | 1.00E+00 | Bacteria | Proteobacteria | Alphaproteobacteria | Rhodobacterales | Rhodobacteraceae | Paracoccus |
| 5634c39a6b4a1475938b6ad7a4113445 | 14.10 | 0.00 | 1.00E+00 | Bacteria | Firmicutes | Bacilli | Lactobacillales | Enterococcaceae | Vagococcus |

baseMean: Mean abundance of the normalized count values over all samples included in the comparison; log2FC: log2-fold change represents the effect size estimate reported on a logarithmic scale to base 2, indicating how much the ASV abundance has changed due to selection for heat-shock resistance in comparison to the control lines; *P* adj: adjusted P-values using the Benjamini-Hochberg procedure.

**Table S7**. Differential abundance analysis comparing lines selected for heat knockdown resistance (KS) to the unselected control lines (UC) at the genus level. Taxonomies were agglomerated to genus level prior to analysis. ASVs are ordered according to adjusted P-values (P adj) from lowest to highest. ASVs were considered differentially abundant for adjusted P-values ≤ 0.01 and an FDR cut-off of 5%. Significance threshold (P adj = 0.01) is marked by a dashed line.

| **ASV ID** | **baseMean** | **log2FC** | ***P* adj** | **Domain** | **Phylum** | **Class** | **Order** | **Family** | **Genus** |
| --- | --- | --- | --- | --- | --- | --- | --- | --- | --- |
| abc8f555ea0b005c9a4085a8343a3f99 | 159.83 | 10.80 | 4.88E-28 | Bacteria | Firmicutes | Bacilli | Bacillales | Paenibacillaceae | Paenibacillus |
| 33b3b41ddb67bb49f482abc6126da323 | 32.73 | 9.59 | 1.19E-19 | Bacteria | Firmicutes | Clostridia | Clostridiales | Lachnospiraceae | Anaerosporobacter |
| 810bf4f2547219682f5ab14a09ca3599 | 2656.56 | -13.04 | 7.91E-17 | Bacteria | Firmicutes | Bacilli | Lactobacillales | Leuconostocaceae | Leuconostoc |
| f8a687d17d5bcd547799c35ad359bf70 | 6790.79 | 7.39 | 1.57E-16 | Bacteria | Proteobacteria | Alphaproteobacteria | Acetobacterales | Acetobacteraceae | Acetobacter |
| 19492e6d689a95489805fe62a81b5ecd | 108.06 | 7.09 | 1.50E-13 | Bacteria | Firmicutes | Bacilli | Bacillales | Paenibacillaceae | Brevibacillus |
| ebc7527f35924613e64e071623af1abb | 53.37 | 9.17 | 3.11E-11 | Bacteria | Firmicutes | Clostridia | Clostridiales | Lachnospiraceae | Anaerocolumna |
| 148a7bc57792072043058510530ea32d | 2331.90 | 5.74 | 9.89E-06 | Bacteria | Firmicutes | Bacilli | Bacillales | Planococcaceae | Lysinibacillus |
| 7406188bce39180c4aa194bfb387935e | 18.48 | 6.49 | 6.27E-05 | Bacteria | Firmicutes | Clostridia | Clostridiales | Lachnospiraceae | Tyzzerella |
| ffaedab25be5e2be043b2355be6644cf | 9.83 | 7.35 | 3.06E-02 | Bacteria | Firmicutes | Clostridia | Clostridiales | Clostridiaceae 1 | Clostridium sensu stricto 1 |
| c37413af5ea0a7254284c92b8bf54ae7 | 225.43 | 2.41 | 4.92E-02 | Bacteria | Proteobacteria | Gammaproteobacteria | Enterobacteriales | Enterobacteriaceae | Providencia |
| f580b2c49c0078358680733c5d838ca6 | 7.02 | 5.75 | 4.99E-02 | Bacteria | Firmicutes | Bacilli | Bacillales | Planococcaceae | Psychrobacillus |
| 19019fa034a9856e8d6365d34574c8fd | 324.39 | -12.61 | 5.32E-02 | Bacteria | Firmicutes | Bacilli | Lactobacillales | Leuconostocaceae | Weissella |
| 05f5c262eb96bf78b6ab2750fdddc0c4 | 18.05 | 3.17 | 8.18E-02 | Bacteria | Firmicutes | Clostridia | Clostridiales | Lachnospiraceae | Lachnoclostridium 5 |
| a406a791970c2c9105ced42674853b40 | 5366.05 | -2.86 | 8.83E-02 | Bacteria | Proteobacteria | Alphaproteobacteria | Rickettsiales | Anaplasmataceae | Wolbachia |
| 9eadd1514e5fcfa10e723634e35f3485 | 197.83 | 1.99 | 2.65E-01 | Bacteria | Actinobacteria | Actinobacteria | Micrococcales | Microbacteriaceae | Microbacterium |
| 4e817d860267844d0bb67bb3ed49522c | 16.36 | -2.31 | 2.65E-01 | Bacteria | Firmicutes | Bacilli | Lactobacillales | Lactobacillaceae | Lactobacillus |
| 0ff98df849b8fae49800a796dadc506c | 49.70 | 0.99 | 5.54E-01 | Bacteria | Firmicutes | Bacilli | Lactobacillales | Enterococcaceae | Enterococcus |
| b9a736421dc15f263c93ec9f509d5f6b | 2.97 | 4.06 | 7.19E-01 | Bacteria | Firmicutes | Clostridia | Clostridiales | Peptostreptococcaceae | Clostridioides |
| 515ecf421a951bc274a7daaa791dc894 | 2.76 | 4.11 | 7.19E-01 | Bacteria | Proteobacteria | Alphaproteobacteria | Rhodobacterales | Rhodobacteraceae | Paracoccus |
| f8d46ee1054ee9224a7e622fd2557cf4 | 3.91 | 1.54 | 9.37E-01 | Bacteria | Firmicutes | Bacilli | Bacillales | Paenibacillaceae | Cohnella |
| 69052d7a94480e1c87649a5aaf431114 | 1.21 | 0.00 | 1.00E+00 | Bacteria | Firmicutes | Clostridia | Clostridiales | Lachnospiraceae | Lachnospiraceae NC2004 group |
| 016b118f0377208f78e5694abe0d59b6 | 2.09 | 0.00 | 1.00E+00 | Bacteria | Firmicutes | Clostridia | Clostridiales | Ruminococcaceae | Ruminiclostridium |
| c2b73b7534124edd282fa67a545e048b | 25.89 | 0.00 | 1.00E+00 | Bacteria | Actinobacteria | Actinobacteria | Corynebacteriales | Nocardiaceae | Rhodococcus |
| cd416ce5bf4afa614154fc0a45feb5a8 | 12.51 | 0.00 | 1.00E+00 | Bacteria | Actinobacteria | Actinobacteria | Micrococcales | Micrococcaceae | Glutamicibacter |
| c63d552898d6b61e6ef07797210d7aca | 14.67 | 0.00 | 1.00E+00 | Bacteria | Actinobacteria | Actinobacteria | Micrococcales | Microbacteriaceae | Leucobacter |
| 43f9d96deeccbd4b7c28e80612ec05ea | 4.96 | 0.17 | 1.00E+00 | Bacteria | Actinobacteria | Actinobacteria | Micrococcales | Brevibacteriaceae | Brevibacterium |
| 7fda73c8ad009ec246fc6a5bf669203e | 860.52 | -0.20 | 1.00E+00 | Bacteria | Actinobacteria | Actinobacteria | Corynebacteriales | Corynebacteriaceae | Corynebacterium 1 |
| b5e95cda7f3db71495303d91e0c9dda7 | 11.99 | 0.00 | 1.00E+00 | Bacteria | Proteobacteria | Gammaproteobacteria | Pseudomonadales | Moraxellaceae | Acinetobacter |
| 64bfc35722d08fe4ccf6b5665b9d1769 | 7.70 | 0.00 | 1.00E+00 | Bacteria | Proteobacteria | Gammaproteobacteria | Enterobacteriales | Enterobacteriaceae | Morganella |
| 5634c39a6b4a1475938b6ad7a4113445 | 14.10 | 0.00 | 1.00E+00 | Bacteria | Firmicutes | Bacilli | Lactobacillales | Enterococcaceae | Vagococcus |

baseMean: Mean abundance of the normalized count values over all samples included in the comparison; log2FC: log2-fold change represents the effect size estimate reported on a logarithmic scale to base 2, indicating how much the ASV abundance has changed due to selection for heat knockdown resistance in comparison to the control lines; *P* adj: adjusted P-values using the Benjamini-Hochberg procedure.

**Table S8**. Differential abundance analysis comparing lines selected for longevity (LS) with unselected control lines (UC) at genus level. Taxonomies were agglomerated to genus level prior to analysis. ASVs are ordered according to adjusted P-values (P adj) from lowest to highest. ASVs were considered differentially abundant for adjusted P-values ≤ 0.01 and an FDR cut-off of 5%. Significance threshold (P adj = 0.01) is marked by a dashed line.

| **ASV ID** | **baseMean** | **log2FC** | ***P* adj** | **Domain** | **Phylum** | **Class** | **Order** | **Family** | **Genus** |
| --- | --- | --- | --- | --- | --- | --- | --- | --- | --- |
| 810bf4f2547219682f5ab14a09ca3599 | 2656.56 | -15.78 | 2.01E-20 | Bacteria | Firmicutes | Bacilli | Lactobacillales | Leuconostocaceae | Leuconostoc |
| abc8f555ea0b005c9a4085a8343a3f99 | 159.83 | 9.09 | 3.83E-20 | Bacteria | Firmicutes | Bacilli | Bacillales | Paenibacillaceae | Paenibacillus |
| 33b3b41ddb67bb49f482abc6126da323 | 32.73 | 8.56 | 6.71E-16 | Bacteria | Firmicutes | Clostridia | Clostridiales | Lachnospiraceae | Anaerosporobacter |
| 19492e6d689a95489805fe62a81b5ecd | 108.06 | 7.39 | 1.24E-14 | Bacteria | Firmicutes | Bacilli | Bacillales | Paenibacillaceae | Brevibacillus |
| f8a687d17d5bcd547799c35ad359bf70 | 6790.79 | 6.20 | 6.22E-12 | Bacteria | Proteobacteria | Alphaproteobacteria | Acetobacterales | Acetobacteraceae | Acetobacter |
| 148a7bc57792072043058510530ea32d | 2331.90 | 8.45 | 1.67E-11 | Bacteria | Firmicutes | Bacilli | Bacillales | Planococcaceae | Lysinibacillus |
| 05f5c262eb96bf78b6ab2750fdddc0c4 | 18.05 | 7.86 | 1.24E-08 | Bacteria | Firmicutes | Clostridia | Clostridiales | Lachnospiraceae | Lachnoclostridium 5 |
| ebc7527f35924613e64e071623af1abb | 53.37 | 7.39 | 1.27E-07 | Bacteria | Firmicutes | Clostridia | Clostridiales | Lachnospiraceae | Anaerocolumna |
| 7406188bce39180c4aa194bfb387935e | 18.48 | 6.44 | 5.73E-05 | Bacteria | Firmicutes | Clostridia | Clostridiales | Lachnospiraceae | Tyzzerella |
| 9eadd1514e5fcfa10e723634e35f3485 | 197.83 | 5.04 | 5.56E-04 | Bacteria | Actinobacteria | Actinobacteria | Micrococcales | Microbacteriaceae | Microbacterium |
| 5634c39a6b4a1475938b6ad7a4113445 | 14.10 | 17.27 | 6.10E-04 | Bacteria | Firmicutes | Bacilli | Lactobacillales | Enterococcaceae | Vagococcus |
| 19019fa034a9856e8d6365d34574c8fd | 324.39 | -13.36 | 3.67E-02 | Bacteria | Firmicutes | Bacilli | Lactobacillales | Leuconostocaceae | Weissella |
| 4e817d860267844d0bb67bb3ed49522c | 16.36 | -3.62 | 4.66E-02 | Bacteria | Firmicutes | Bacilli | Lactobacillales | Lactobacillaceae | Lactobacillus |
| cd416ce5bf4afa614154fc0a45feb5a8 | 12.51 | 5.28 | 5.37E-02 | Bacteria | Actinobacteria | Actinobacteria | Micrococcales | Micrococcaceae | Glutamicibacter |
| c37413af5ea0a7254284c92b8bf54ae7 | 225.43 | 2.13 | 6.79E-02 | Bacteria | Proteobacteria | Gammaproteobacteria | Enterobacteriales | Enterobacteriaceae | Providencia |
| ffaedab25be5e2be043b2355be6644cf | 9.83 | 5.72 | 8.04E-02 | Bacteria | Firmicutes | Clostridia | Clostridiales | Clostridiaceae 1 | Clostridium sensu stricto 1 |
| 7fda73c8ad009ec246fc6a5bf669203e | 860.52 | -1.25 | 1.82E-01 | Bacteria | Actinobacteria | Actinobacteria | Corynebacteriales | Corynebacteriaceae | Corynebacterium 1 |
| a406a791970c2c9105ced42674853b40 | 5366.05 | -2.28 | 1.82E-01 | Bacteria | Proteobacteria | Alphaproteobacteria | Rickettsiales | Anaplasmataceae | Wolbachia |
| f8d46ee1054ee9224a7e622fd2557cf4 | 3.91 | 4.42 | 2.32E-01 | Bacteria | Firmicutes | Bacilli | Bacillales | Paenibacillaceae | Cohnella |
| 515ecf421a951bc274a7daaa791dc894 | 2.76 | 6.18 | 3.87E-01 | Bacteria | Proteobacteria | Alphaproteobacteria | Rhodobacterales | Rhodobacteraceae | Paracoccus |
| f580b2c49c0078358680733c5d838ca6 | 7.02 | 1.96 | 6.27E-01 | Bacteria | Firmicutes | Bacilli | Bacillales | Planococcaceae | Psychrobacillus |
| 0ff98df849b8fae49800a796dadc506c | 49.70 | 0.56 | 7.77E-01 | Bacteria | Firmicutes | Bacilli | Lactobacillales | Enterococcaceae | Enterococcus |
| 016b118f0377208f78e5694abe0d59b6 | 2.09 | 1.86 | 9.59E-01 | Bacteria | Firmicutes | Clostridia | Clostridiales | Ruminococcaceae | Ruminiclostridium |
| 69052d7a94480e1c87649a5aaf431114 | 1.21 | 0.00 | 1.00E+00 | Bacteria | Firmicutes | Clostridia | Clostridiales | Lachnospiraceae | Lachnospiraceae NC2004 group |
| b9a736421dc15f263c93ec9f509d5f6b | 2.97 | 0.00 | 1.00E+00 | Bacteria | Firmicutes | Clostridia | Clostridiales | Peptostreptococcaceae | Clostridioides |
| c2b73b7534124edd282fa67a545e048b | 25.89 | 0.00 | 1.00E+00 | Bacteria | Actinobacteria | Actinobacteria | Corynebacteriales | Nocardiaceae | Rhodococcus |
| c63d552898d6b61e6ef07797210d7aca | 14.67 | 0.00 | 1.00E+00 | Bacteria | Actinobacteria | Actinobacteria | Micrococcales | Microbacteriaceae | Leucobacter |
| 43f9d96deeccbd4b7c28e80612ec05ea | 4.96 | -0.31 | 1.00E+00 | Bacteria | Actinobacteria | Actinobacteria | Micrococcales | Brevibacteriaceae | Brevibacterium |
| b5e95cda7f3db71495303d91e0c9dda7 | 11.99 | 0.00 | 1.00E+00 | Bacteria | Proteobacteria | Gammaproteobacteria | Pseudomonadales | Moraxellaceae | Acinetobacter |
| 64bfc35722d08fe4ccf6b5665b9d1769 | 7.70 | 0.00 | 1.00E+00 | Bacteria | Proteobacteria | Gammaproteobacteria | Enterobacteriales | Enterobacteriaceae | Morganella |

baseMean: Mean abundance of the normalized count values over all samples included in the comparison; log2FC: log2-fold change represents the effect size estimate reported on a logarithmic scale to base 2, indicating how much the ASV abundance has changed due to selection for longevity in comparison to the control lines; P adj: adjusted P-values using the Benjamini-Hochberg procedure.

**Table S9**. Differential abundance analysis comparing lines selected for starvation resistance (SS) to the unselected control lines (UC) at the genus level. Taxonomies were agglomerated to genus level prior to analysis. ASVs are ordered according to adjusted P-values (P adj) from lowest to highest. ASVs were considered differentially abundant for adjusted P-values ≤ 0.01 and an FDR cut-off of 5%. Significance threshold (P adj = 0.01) is marked by a dashed line.

| **ASV ID** | **BaseMean** | **log2FC** | ***P* adj** | **Domain** | **Phylum** | **Class** | **Order** | **Family** | **Genus** |
| --- | --- | --- | --- | --- | --- | --- | --- | --- | --- |
| abc8f555ea0b005c9a4085a8343a3f99 | 159.83 | 10.08 | 1.63E-24 | Bacteria | Firmicutes | Bacilli | Bacillales | Paenibacillaceae | Paenibacillus |
| 33b3b41ddb67bb49f482abc6126da323 | 32.73 | 7.03 | 1.10E-10 | Bacteria | Firmicutes | Clostridia | Clostridiales | Lachnospiraceae | Anaerosporobacter |
| 810bf4f2547219682f5ab14a09ca3599 | 2656.56 | -8.71 | 2.50E-09 | Bacteria | Firmicutes | Bacilli | Lactobacillales | Leuconostocaceae | Leuconostoc |
| ebc7527f35924613e64e071623af1abb | 53.37 | 8.02 | 1.34E-08 | Bacteria | Firmicutes | Clostridia | Clostridiales | Lachnospiraceae | Anaerocolumna |
| 05f5c262eb96bf78b6ab2750fdddc0c4 | 18.05 | 7.42 | 1.17E-07 | Bacteria | Firmicutes | Clostridia | Clostridiales | Lachnospiraceae | Lachnoclostridium 5 |
| c37413af5ea0a7254284c92b8bf54ae7 | 225.43 | 5.46 | 2.31E-07 | Bacteria | Proteobacteria | Gammaproteobacteria | Enterobacteriales | Enterobacteriaceae | Providencia |
| 5634c39a6b4a1475938b6ad7a4113445 | 14.10 | 22.78 | 3.84E-06 | Bacteria | Firmicutes | Bacilli | Lactobacillales | Enterococcaceae | Vagococcus |
| 7406188bce39180c4aa194bfb387935e | 18.48 | 7.23 | 4.54E-06 | Bacteria | Firmicutes | Clostridia | Clostridiales | Lachnospiraceae | Tyzzerella |
| 19492e6d689a95489805fe62a81b5ecd | 108.06 | 4.23 | 1.98E-05 | Bacteria | Firmicutes | Bacilli | Bacillales | Paenibacillaceae | Brevibacillus |
| c2b73b7534124edd282fa67a545e048b | 25.89 | 8.88 | 3.74E-04 | Bacteria | Actinobacteria | Actinobacteria | Corynebacteriales | Nocardiaceae | Rhodococcus |
| c63d552898d6b61e6ef07797210d7aca | 14.67 | 8.03 | 2.15E-03 | Bacteria | Actinobacteria | Actinobacteria | Micrococcales | Microbacteriaceae | Leucobacter |
| 9eadd1514e5fcfa10e723634e35f3485 | 197.83 | 4.26 | 3.96E-03 | Bacteria | Actinobacteria | Actinobacteria | Micrococcales | Microbacteriaceae | Microbacterium |
| cd416ce5bf4afa614154fc0a45feb5a8 | 12.51 | 7.35 | 3.96E-03 | Bacteria | Actinobacteria | Actinobacteria | Micrococcales | Micrococcaceae | Glutamicibacter |
| 148a7bc57792072043058510530ea32d | 2331.90 | 3.60 | 6.62E-03 | Bacteria | Firmicutes | Bacilli | Bacillales | Planococcaceae | Lysinibacillus |
| 4e817d860267844d0bb67bb3ed49522c | 16.36 | -4.23 | 1.26E-02 | Bacteria | Firmicutes | Bacilli | Lactobacillales | Lactobacillaceae | Lactobacillus |
| 19019fa034a9856e8d6365d34574c8fd | 324.39 | -14.49 | 1.53E-02 | Bacteria | Firmicutes | Bacilli | Lactobacillales | Leuconostocaceae | Weissella |
| f8a687d17d5bcd547799c35ad359bf70 | 6790.79 | -2.11 | 2.82E-02 | Bacteria | Proteobacteria | Alphaproteobacteria | Acetobacterales | Acetobacteraceae | Acetobacter |
| f580b2c49c0078358680733c5d838ca6 | 7.02 | 5.54 | 3.70E-02 | Bacteria | Firmicutes | Bacilli | Bacillales | Planococcaceae | Psychrobacillus |
| b5e95cda7f3db71495303d91e0c9dda7 | 11.99 | 8.54 | 8.50E-02 | Bacteria | Proteobacteria | Gammaproteobacteria | Pseudomonadales | Moraxellaceae | Acinetobacter |
| 43f9d96deeccbd4b7c28e80612ec05ea | 4.96 | -3.92 | 2.28E-01 | Bacteria | Actinobacteria | Actinobacteria | Micrococcales | Brevibacteriaceae | Brevibacterium |
| ffaedab25be5e2be043b2355be6644cf | 9.83 | 3.75 | 2.65E-01 | Bacteria | Firmicutes | Clostridia | Clostridiales | Clostridiaceae 1 | Clostridium sensu stricto 1 |
| 69052d7a94480e1c87649a5aaf431114 | 1.21 | 4.52 | 4.33E-01 | Bacteria | Firmicutes | Clostridia | Clostridiales | Lachnospiraceae | Lachnospiraceae NC2004 group |
| 0ff98df849b8fae49800a796dadc506c | 49.70 | 0.88 | 4.76E-01 | Bacteria | Firmicutes | Bacilli | Lactobacillales | Enterococcaceae | Enterococcus |
| f8d46ee1054ee9224a7e622fd2557cf4 | 3.91 | 2.47 | 5.26E-01 | Bacteria | Firmicutes | Bacilli | Bacillales | Paenibacillaceae | Cohnella |
| 7fda73c8ad009ec246fc6a5bf669203e | 860.52 | 0.19 | 9.36E-01 | Bacteria | Actinobacteria | Actinobacteria | Corynebacteriales | Corynebacteriaceae | Corynebacterium 1 |
| a406a791970c2c9105ced42674853b40 | 5366.05 | 0.34 | 9.36E-01 | Bacteria | Proteobacteria | Alphaproteobacteria | Rickettsiales | Anaplasmataceae | Wolbachia |
| 64bfc35722d08fe4ccf6b5665b9d1769 | 7.70 | -0.37 | 9.76E-01 | Bacteria | Proteobacteria | Gammaproteobacteria | Enterobacteriales | Enterobacteriaceae | Morganella |
| b9a736421dc15f263c93ec9f509d5f6b | 2.97 | 0.00 | 1.00E+00 | Bacteria | Firmicutes | Clostridia | Clostridiales | Peptostreptococcaceae | Clostridioides |
| 016b118f0377208f78e5694abe0d59b6 | 2.09 | 0.00 | 1.00E+00 | Bacteria | Firmicutes | Clostridia | Clostridiales | Ruminococcaceae | Ruminiclostridium |
| 515ecf421a951bc274a7daaa791dc894 | 2.76 | 0.00 | 1.00E+00 | Bacteria | Proteobacteria | Alphaproteobacteria | Rhodobacterales | Rhodobacteraceae | Paracoccus |

baseMean: Mean abundance of the normalized count values over all samples included in the comparison; log2FC: log2-fold change represents the effect size estimate reported on a logarithmic scale to base 2, indicating how much the ASV abundance has changed due to selection for starvation resistance in comparison to the control lines; *P* adj: adjusted P-values using the Benjamini-Hochberg procedure.
